# Short Term Impact of Recycling-Derived Fertilizers on Their P Supply for Perennial Ryegrass (*Lolium perenne*)

**DOI:** 10.1101/2023.03.29.534721

**Authors:** Lea Deinert, Bastian Egeter, Israel Ikoyi, Patrick Forrestal, Achim Schmalenberger

## Abstract

Phosphorus is a finite, essential macronutrient for agriculture. Various nutrient recycling technologies in waste streams management are currently under development in many European countries in order to alleviate the dependency of the EU on imports of non-renewable raw material for the production of mineral phosphorus fertilizers commonly used in agriculture. The resulting products such as struvites and ashes need to be assessed for their application as so-called recycling-derived fertilisers (RDF) in the agricultural sector prior to commercialisation. Albeit high phosphorus abundance in most soils, the phosphorus availability for plant growth promotion in the soil solution is usually low due to strong P sorption in soil and depends vastly on the microbial mobilisation capability of the soil.

To investigate the impact of different phosphorus fertilizers on plant growth and the soil P cycling microbiota, a short-term pot trial was conducted over the period of 54 days. *Lolium perenne* (var. AberGreen) was grown with application of superphosphate (SP) as inorganic fertiliser, two ashes (poultry litter ash (PLA) and sewage sludge ash (SSA) and two struvites (municipal wastewater struvite (MWS) and commercial CrystalGreen^®^ (CGS) in rates of 20 and 60 kg P ha^-1^ in four replicates. A P-free control (SP0) was also included in the trial.

Post-harvest, a positive correlation between dry weight yield and struvite application was detected, struvite P also was higher readily available and ACP activity was significantly improved for struvites at the high P application rate. The ash RDFs showed a liming effect at 60 kg P ha^-1^, and PLA60 negatively affected ACP activity, while PLA20 had significantly lower *phoD* copy numbers. P mobilization from phosphonates and phytates was not affected, TCP solubilization was negatively affected by mineral SP fertilizer application at both P concentrations. Overall, the bacterial and *phoD* harbouring community were not strongly affected by the P fertilization in this study.

## Introduction

Phosphorus (P) is present in all life, such as a part of the energy-carrying molecule ATP, and DNA and RNA (Elser, 2012). P is an essential macronutrient for plants and is therefore a crucial element for food production. Intensification of modern agriculture requires P fertilizer application to soil to compensate for the P off-take by plant harvests and retain soil productivity (Roy *et al*., 2016). Current P management is facilitated in a linear manner, where P is extracted as phosphate rock from mines located outside Europe with the largest abundance in Morocco and the Western Sahara, processed to mineral P fertilizers and applied to land. The excessive use of these fertilizers to improve crop productivity causes P run-off and leaching, one of the main causes of eutrophication of water bodies (Bindraban *et al*., 2020). Additionally, the handling of P-rich wastes, either from food production/agriculture, wastewater or industrial wastes are not optimally managed. The focus has been mostly on P removal than P recovery due to measures to limit environmental pollution, however, these practices need to shift towards a more sustainable approach (Commission, 1991, Carpenter & Bennett, 2011). P cannot be substituted by any other element, and it is becoming increasingly scarce. Its depletion as fossil resource, the main source for mineral P fertilizer production, has been predicted within the next few centuries (van Kauwenbergh, 2010, Cordell *et al*., 2011).

Another issue is the negative impact of mineral P fertilizer on the soil fertility in general. While fertilizers enhance crop production in the short term, their impact on the soil P cycle and the ability of the soil microbiota to replenish P in the long term can be seriously affected. The soil microbiota, constituting of a plethora of bacteria and fungi, has an important role in nutrient cycling and overall soil structure and fertility (Richardson & Simpson, 2011, Bano & Iqbal, 2016). Soil microbes are involved in organic matter decomposition, nutrient mineralisation and immobilization, and interactions between bacteria, fungi and plants can be established to facilitate nutrient acquisition (Custódio *et al*., 2022). Increased microbial activity is usually witnessed in the soil rhizosphere, which describes the soil volume influenced by the plant root. During uptake of bioavailable orthophosphate by plant roots, a P depletion zone is created. The plants exude a range of compounds, such as amino acids, organic acids, sugars and vitamins, which influence the soil microbes habituating the soil rhizosphere, stimulating microbial nutrient mobilization activities from P pools that are inaccessible for plants (Mukerji *et al*., 2006). Reductions in soil biodiversity and therefore a loss in functionality of the soil nutrient cycle have been reported upon long-term mineral P fertilizer application and change of land use (McLaughlin & Mineau, 1995, Mozumder & Berrens, 2007, Kaminsky *et al*., 2018, Borrelli *et al*., 2020, Liu *et al*., 2020).

In order to establish global food security and to avoid or reduce dependence upon unbalanced distributed resources located in geopolitically unstable countries and ecological implications of environmental nutrient pollution, nutrient recovery from waste streams has been investigated in more detail in recent years (Cornel & Schaum, 2009, Roy, 2017). Potential sources for P recovery include wastewater, sewage sludge and manure. The resulting products of a multitude of P recovery processes are termed recycling-derived fertilizers (RDFs) or bio-based fertilizers (Chojnacka *et al*., 2020, Postma *et al*., 2020). While the manufacturing units used to produce RDFs are mostly at pilot-production scale, which makes their current production costly and non-competitive compared to conventional mineral P fertilizers, these technologies might gain increased importance in the near future, when high P demand or military conflicts in producer countries causes fertilizer price surges (Mogollón *et al*., 2018, El Wali *et al*., 2019). Furthermore, it was reported that sewage sludge disposal accounted for 40% of the greenhouse gas emissions of wastewater treatment plants. This could be significantly reduced under adoption of a circular economy and enable sludge valorisation for RDF production (Brown *et al*., 2010, Gherghel *et al*., 2019). According Ramankutty and colleagues (Ramankutty *et al*., 2008), 70% of agriculturally used land in the world comprises grassland, in Ireland even around 90% of its agriculture is dominated by grass-based systems (Virto *et al*., 2014). Therefore, evaluation of these novel fertilizer products as alternative to phosphate rock-based P fertilizers in grasslands is beneficial to support their widespread use in these agroecological systems.

In this experiment, four RDFs were evaluated regarding their P supply for perennial ryegrass (*Lolium perenne*) cultivation in comparison to superphosphate in pots. Two different P concentrations of 20 and 60 kg P ha^-1^ were used for each fertilizer, and all fertilized treatments were compared to a P-free control. It was hypothesized that a) the RDFs would have similar fertilization value as conventional superphosphate at low and high P fertilization and b) RDFs have a lower impact on the soil microbiota involved in P cycling. The objective was to analyse the abundance, composition and function of the soil P mobilizing bacteria under different P fertilizer products as well as no P addition and the resulting impact on grassland productivity.

## Materials and Methods

### Pot Trial Set-Up and Fertilizer Preparation

The soil used in this experiment was a sandy loam soil (51% sand, 42% silt, 7% clay) with a pH (measured in 0.01 M CaCl_2_) of 5.0, a cation exchange capacity of 10.5 meq 100 g^-1^, organic matter content (determined via loss-on-ignition) of 5.3% and a Morgan’s P index of 2 (4.2 mg P L^-1^), indicating a likely crop response to the fertilizer application. The soil was taken from a field at Teagasc Johnstown Castle (Wexford, Ireland, N52°17’47”, W6°30’29”), which had been reseeded with perennial ryegrass in 2018 and prior to that was under long-term pasture management (personal communication with Patrick Forrestal). The soil was air-dried to a residual moisture content of 10 – 20% to allow for mixing, homogenizing, and sieving of the soil on one hand and to prevent damages to the soil microbes via complete dehydration on the other hand. Then, the soil was sieved through a sieve with a mesh size of 5.60 mm, to exclude bigger stones, followed by passing the soil through a smaller sieve (mesh size 3.35 mm) to remove smaller gravel, plant roots and to break down bigger soil aggregates. The moisture content and field capacity (water holding capacity) of the sieved soil was then determined. Eleven treatments were prepared in quadruplicates, in detail the RDFs poultry litter ash (PLA), sewage sludge ash (SSA), commercially available struvite (CrystalGreen^®^, CGS) and struvite from a municipal wastewater plant (MWS) and a P-free control (SP0) were used. The suitability of the RDFs was compared to a superphosphate (SP) fertilizer (SP40), which is a mixture of triple superphosphate and a filler (ratio 80:20) containing 16% P and 15% Ca and is commonly applied in grasslands in Ireland (personal communication with Patrick Forrestal). The SP and the struvite fertilizers exhibited different granule sizes, with SP and CGS possessing large granules and MWS has small ones, as illustrated in Figure 1.

**Figure 1.**
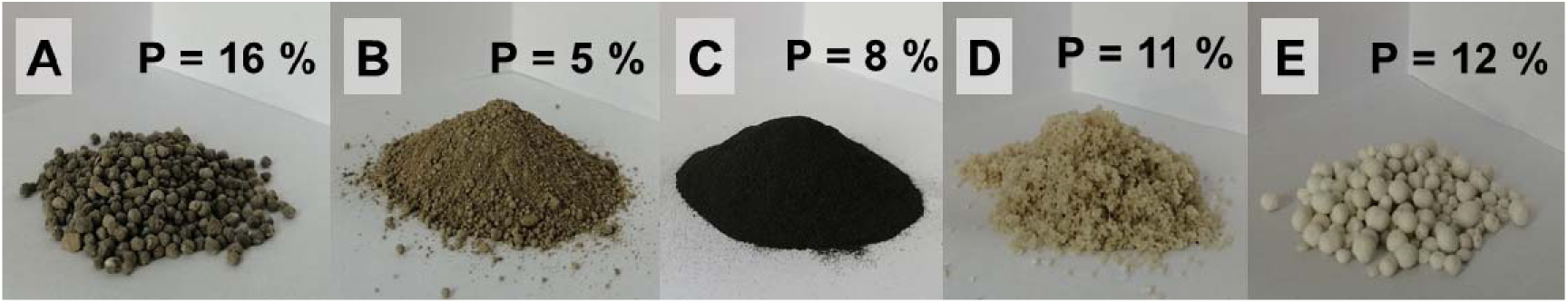
Images of P fertilizers applied in the pot trial with their P content given in percent: A) superphosphate (SP), followed by recycling-derived fertilizers B) poultry litter ash (PLA), C) sewage sludge ash (SSA), D) municipal wastewater struvite (MWS) and E) CrystalGreen^®^ struvite (CGS).

Therefore, the larger granules were broken down (not ground) with a pestle and mortar to obtain smaller pieces of approximately 1mm in diameter, and therefore potentially reducing bias by achieving higher accuracy in weighing of the products and improved similarity of distribution of the fertilizers, when mixed with soil. Moreover, the MWS fertilizer appeared to be moist and was therefore dried in a fan oven at 40°C, until a weight equilibrium had been reached. The drying temperature for struvites did not exceed 40°C to avoid N losses. P fertilization was carried out at two different concentrations: 20 kg ha^-1^, which is typical for grassland fertilization and also used for P build-up in soil and 60 kg ha^-1^, which is the recommended amount for pasture establishment (Plunkett *et al*., 2016). The nutritional composition of the RDFs is displayed in Table 1.

**Table 1.** Nutritional composition of the recycling-derived fertilizers applied in the experiment, the values for the RDFs PLA, SSA and MWS were evaluated by University of Ghent (ReNu2Farm report, WPT1 D3.1. Product Characterization) and the values for CGS were given on the company’s website (https://crystalgreen.com/agriculture/).

|  | PLA | SSA | MWS | CGS |
| --- | --- | --- | --- | --- |
| pH (KCl) | 12.0 | 11.2 | 7.1 | n.d. |
| Dry matter (%) | 92.0 | 100.0 | 50.0 | n.d. |
| Total Carbon (g kg<sup>-1</sup> FW) | 10.7 | < 5.5 | < 5.5 | n.d. |
| Total Nitrogen (g kg<sup>-1</sup> FW) | <0.9 | <0.9 | 53.1 | n.d. |
| NH<sub>4</sub>-N (g kg<sup>-1</sup> FW) | 0.01 | 0.02 | 0.11 | 50 |
| P (g kg<sup>-1</sup> FW) | 38.7 | 48.2 | 49.8 | 120 |
| K (g kg<sup>-1</sup> FW) | 69.8 | 6.7 | 0.21 | n.d. |
| S (g kg<sup>-1</sup> FW) | 19.2 | 39.9 | 0.04 | n.d. |
| Na (g kg<sup>-1</sup> FW) | 9.1 | 104.6 | 0.03 | n.d. |
| Ca (g kg<sup>-1</sup> FW) | 111.9 | 62.7 | 0.18 | n.d. |
| Mg (g kg<sup>-1</sup> FW) | 26.5 | 8.9 | 44.3 | 100 |
| Zn (mg kg<sup>-1</sup> FW) | 1592.9 | 1390.3 | < 4.17 | n.d. |
| Fe (mg kg<sup>-1</sup> FW) | 3708.8 | 31939.5 | 118.8 | n.d. |
| Cu (mg kg<sup>-1</sup> FW) | 312.5 | 460.5 | < 3.33 | n.d. |
| Al (mg kg<sup>-1</sup> FW) | 4972.0 | 35541.6 | < 6.67 | n.d. |

FW: fresh weight, n.d.: not determined

All fertilizers were applied to the soil in advance of setting up the pots and the soil was thoroughly mixed upon fertilizer addition to avoid differences in micro-environmental conditions in the short-term trial. Additionally, because the amount of fertilizer was less than 0.5 g for the low P rate and less than 1.3 g for the high P rate per pot, the mixing of soil and fertilizer was carried out for each pot individually to ensure that every pot received the same amount of fertilizer. The mixture was applied in equal amounts of 825 g to the round plant pots (height = 10 cm, diameter_top_ = 12 cm, diameter_bottom_ = 9 cm), after a nylon mesh (20 µm mesh size, PlastOK, Birkenhead, UK) was added to the bottom of the pot to minimize root growth outside the pot and reduce the loss of bulk soil from the pot. On top of the soil and fertilizer mix, a small layer of untreated soil was applied (approx. 3cm) and the *Lolium perenne* (diploid var. AberGreen, provided by Germinal Ireland Ltd., Tipperary, Ireland) seeds were added to allow for similar seed germination conditions in all treatments without fertilizer contact. The amount of seeds applied was derived from sowing recommendations received from Teagasc (Agricultural and Food Development Authority in Ireland) for a seed mixture of diploid (D) and tetraploid (T) varieties of *Lolium perenne* (40 % AberGreen (D), 30 % AberChoice (D) and 30 % AberGain (T)) at a rate of 14 lb ac^-1^ (34.6 kg ha^-1^). The weight difference between the diploid and tetraploid seed mixture was corrected by counting and weighing 100 seeds of both the mixed varieties and the single variety in triplicates and using the weight ratio. Furthermore, the amount of seeds was adjusted for the low germination rate of 17% in soil tested beforehand. The calculation of the seed amount to be weighted in for each pot with all considerations mentioned above was carried out as follows:

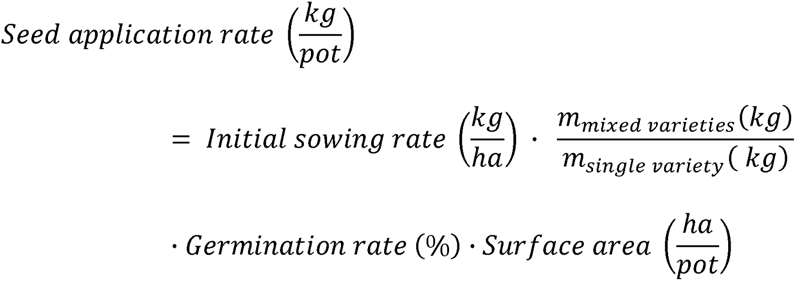

Finally, the pots were watered to obtain 70% of the water holding capacity of the soil. The pots were placed on pot saucers with drains to prevent stagnant moisture. The trial was run for 54 days in a greenhouse. Irrigation of pots was carried out with rainwater collected at the Field Biological Unit at the University of Limerick every second day with an initial amount of 15 mL of water, which was gradually increased to 25 mL to account for plant needs. Two samples of the rainwater were taken throughout the course of the experiment to determine the nutrient composition and to confirm, that the P intake from this source was negligible. The pots were weighed back once a week to monitor the moisture status.

Shortly after seed germination, a full complement of the nutrients nitrogen (N), potassium (K), sulphur (S), magnesium (Mg), calcium (Ca), zinc (Zn) and copper (Cu) was added to prevent starvation effects in the ryegrass due to lack of any other nutrient than P. The nutrient requirements for perennial ryegrass were obtained from Teagasc (Plunkett and Wall 2016), as shown in Table 2.

**Table 2.** Nutrient application rates as recommended by Teagasc for grassland on which grazing livestock is kept, and the nutrient concentrations finally applied to the pots (Plunkett & Wall, 2016).

|  | Recommended<br>Concentrations<br>(kg ha<sup>-1</sup>) | Concentration<br>Applied<br>(kg ha<sup>-1</sup>) | Concentration<br>(mg cm<sup>-2</sup>) | Mass per Pot<br>(mg) |
| --- | --- | --- | --- | --- |
| N | 220 | 220 | 2.20 | 248.8 |
| K | 182 | 185 | 1.85 | 209.2 |

|  | **Recommended**<br>**Concentrations**<br>**(kg ha<sup>-1</sup>)** | **Concentration**<br>**Applied**<br>**(kg ha<sup>-1</sup>)** | **Concentration**<br>**(mg cm<sup>-2</sup>)** | **Mass per Pot**<br>**(mg)** |
| --- | --- | --- | --- | --- |
| Ca | 2 kg Ca / 1 kg N | 440 | 4.40 | 497.6 |
| S | 20 | 20 | 0.20 | 22.6 |
| Mg | 30 | 30 | 0.12 | 13.7 |
| ZnSO<sub>4</sub> | 5 | 5 | 0.05 | 5.7 |
| CuSO<sub>4</sub> | 20 | 15 | 0.15 | 17.0 |

The application of the nutrients in the form of solutions was conducted on two subsequent irrigation days, instead of adding water. However, prior to the fertilizer solution application, 5 mL of water were added to precondition the soil. The solutions prepared from KNO_3_, Ca(NO_3_)_2_, MgCl_2_ and MgSO_4_ were added on the first occasion and CaCl_2_, CuSO_4_ and ZnSO_4_ were supplemented on the following irrigation day.

### Harvest Procedure of Plants and Soil

The experiment was terminated after 54 days of perennial ryegrass cultivation. The pot irrigation was stopped five days in advance of the harvest to ensure that the soil was not too wet, so that separation of bulk and rhizosphere soil was successful during the disassembly of the pots. An overview of the experimental workflow can be found in Figure 2. The bulk soil of each pot was collected in one large bag per treatment by gently shaking off loosely attached soil after removal of the roots and soil from the pot. Then the roots were put in zip-lock bags and the rhizosphere soil, which is closely attached to the plant roots was shaken off more vigorously. The rhizosphere soil was stored at 4°C immediately after harvest. Aliquots were taken for storage at −20°C for DNA extraction at a later stage. The plant shoots were then cut off the roots, the shoots were placed in paper bags to determine the fresh weight per pot and the dry weight after drying for 72 h at 55°C. Elemental analysis of the dried shoots was performed at Lancrop Laboratories Ltd., where atomic absorption spectroscopy, inductively coupled plasma spectrometry, spectrophotometry and titrations were employed according to ISO/IEC 17025:2005 to determine the boron, calcium, copper, iron, manganese, magnesium, molybdenum, nitrogen, phosphorus, potassium, sulphur and zinc content per pooled treatment.

**Figure 2.**
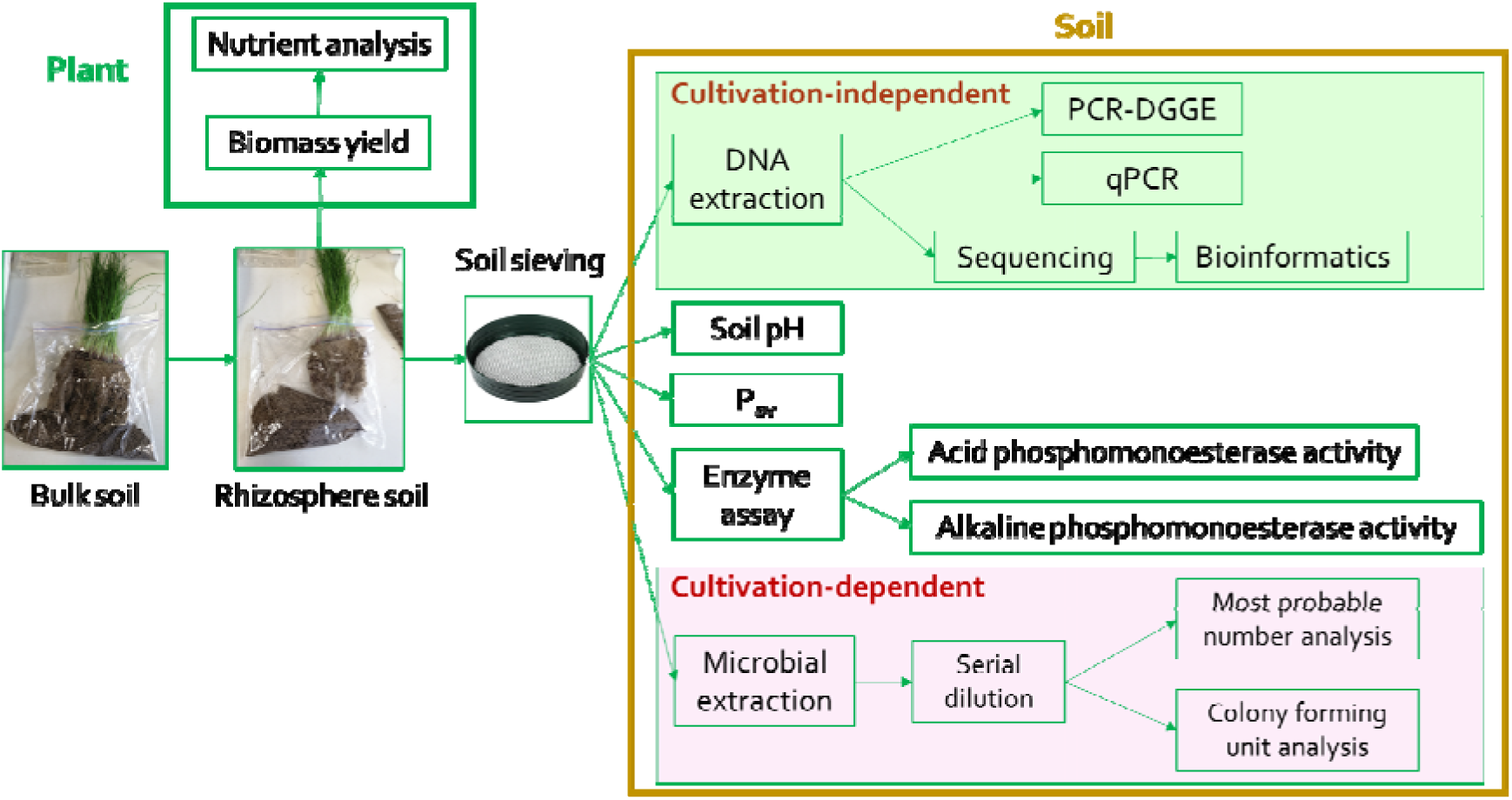
Experimental workflow of the pot trial post-harvest analyses.

**Figure 3.**
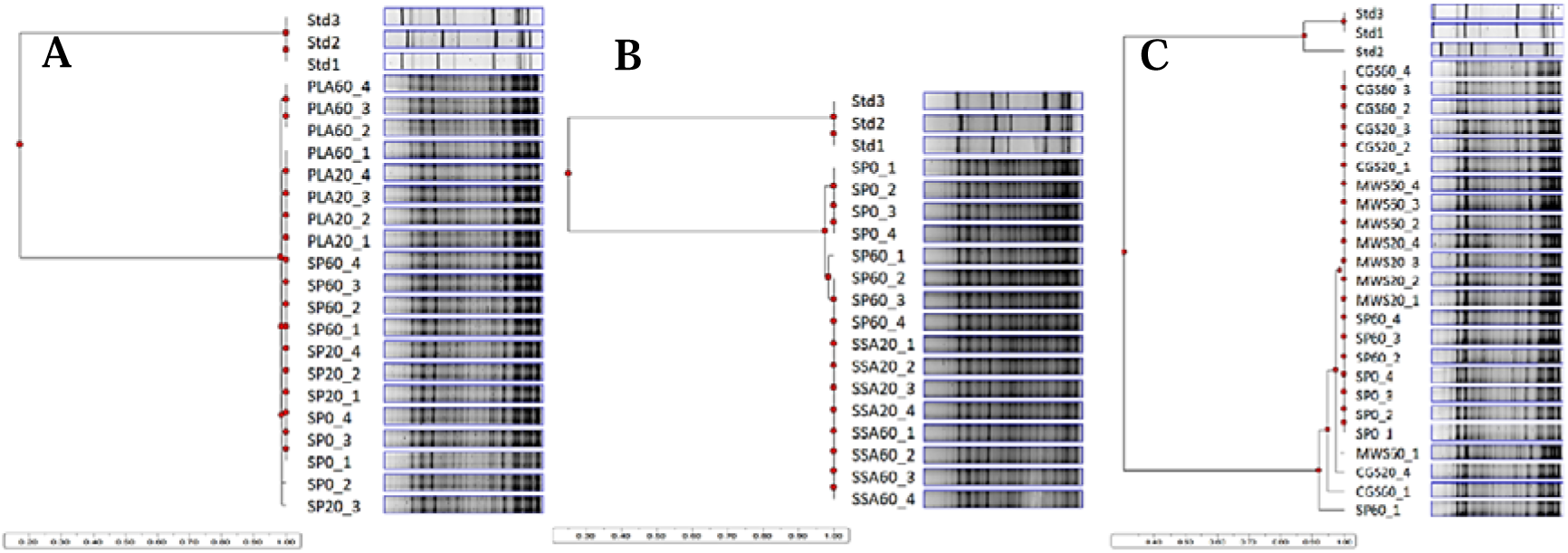
UPGMA dendrograms for all 16S rRNA PCR-DGGEs performed after DNA extraction from the pot trial rhizosphere soil, A comparison of the band patterns between treatments SP0, SP20, SP60 and the RDFs PLA20 and PLA60, B comparison between SP0, SP60 and the RDFs SSA20 and SSA60, C comparison between SP0, SP60 and RDFs MWS20, MWS60, CGS20 and CGS60, n=4.

### Post-Harvest Experiments

The bulk soil was used to determine the soil pH for each sample. As stated previously (Fox *et al*., 2014) 5 g of air-dried and sieved soil were suspended in 20 mL of 0.01 M CaCl_2_ solution and rotated at 70 rpm for 5 min on an end-over-end RS-2M Intellimixer (ELMI, Riga, Latvia). After allowing the solution to settle for 2 h, the pH was measured potentiometrically in the supernatant. All other experiments were carried out with the rhizosphere soil.

For the MPN approach and the CFU analysis, a serial dilution of bacterial extracts from rhizosphere soil was carried out, as described previously (Fox *et al*., 2014). There, 1 g of rhizosphere soil was weighed into a 50 mL assay tube. 10 mL of sterile saline solution (0.85% w/v NaCl, stored at 4°C) were added aseptically and the suspension was rotated at 75 rpm for 30 min at 4°C. 0.1 mL was taken immediately before the solution started settling and was applied seven times in a ten-fold serial dilution in 0.9 mL sterile saline solution in a laminar flow cabinet. Microtiter plates were filled with 180 µL MM2PAA medium, containing only phosphonate as single P source, MM2Phy medium, which contains only phytate (inositol hexaphosphoric acid) as single P source or R2 medium, a nutrient reduced medium for heterotrophic growth (Reasoner & Geldreich, 1985, Fox *et al*., 2014). Then, 20 µL of each dilution step (10^-1^ to 10^-7^) was applied in 5 technical replicates to the plates. The plates were closed with a lid, wrapped in parafilm to prevent evaporation, and incubated at 25°C for 14 days. After 14 days of incubation the microtiter plate wells were assessed for cloudiness or colour change, indicating microbial growth. The number of positive wells, where growth had been observed, per dilution step were recorded in a table and a three-digit MPN value was derived using the dilution step with the highest dilution, where all wells were recorded as positive, and the number of positive wells of the two following higher dilutions. This MPN value was then compared to a MPN table provided by the Food and Drug Administration (FDA, Blodgett, https://www.fda.gov/food/laboratory-methods-food/bam-appendix-2-most-probable-number-serial-dilutions) to obtain the MPN g^-1^ value corresponding to the three-digit code. This value must be multiplied with the dilution factor of the highest dilution, where all wells were positive for microbial growth, then the dilution factor for the addition to the well (50 μL mL^-1^), the dilution factor for the initial soil sample amount (in grams) applied in saline solution and the final result was expressed as MPN g soil^-1^, always in conjunction with the P source that had been applied to the minimal medium.

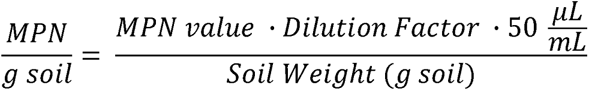

100 μL of selected dilutions (10^-3^ and 10^-4^) were applied to an individual TCP plate (Fox *et al*., 2014) in three technical replicates and distributed on the surface with a sterile (Drigalski) spreader, until the liquid has been absorbed by the agar. The plates were then sealed with parafilm and were incubated at 25°C in the dark for 14 days. The cultivation of soil microbes on the solid medium supplemented with tri-calcium phosphate was analysed by counting the colonies which formed a halo around them, a clearing zone indicating the solubilization of the TCP in the agar. The count was then multiplied with the dilution factor of the inoculation, corrected for the volume from 100 µL to 1 mL and adjusted for the soil weight used for the microbial extraction:

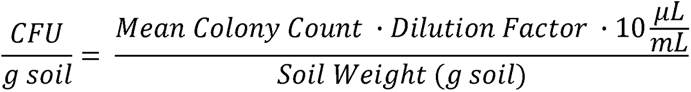

Measuring the potential acid and alkaline phosphomonoesterase activity in soil was conducted following standard protocols (Tabatabai & Bremner, 1969). First, 1 g of soil (sieved, 2 mm mesh size) was mixed with 0.2 mL toluene, 4 mL modified universal buffer (either at pH 6.5 for acid phosphatase, or at pH 11 for alkaline phosphatase) and 1 mL 0.05 M p-nitrophenyl phosphate (pNP) as substrate for the hydrolytic reaction and then incubated in a water bath at 37°C for an hour. After the incubation had finished, 1 mL 0.5 M calcium chloride and 4 mL 0.5 M sodium hydroxide were added, and the solution was mixed to stop the enzyme reaction. Controls were prepared similarly as a composite sample for each treatment, however, the p-nitrophenyl phosphate solution was added after the enzymatic reaction had been stopped. This was done to correct for background signal originating from other sources than p-nitrophenol release in the sample. The soil suspension was then filtered through Grade 113 Cellulose qualitative filter paper (Fisherbrand, Fisher Scientific, UK) and the resulting p-nitrophenol was measured spectrophotometrically (M501 Single Beam Scanning UV/Visible Spectrophotometer, Camspec, UK) at 420 nm. The final p-nitrophenol concentration obtained in 1 g of soil after 1 h of substrate incubation was calculated via a standard curve by measuring the absorption for 0 (blank), 10, 20, 30, 40 and 50 µg p-nitrophenol and the result was stated in μg g soil^-1^ h^-1^.

The available P was measured using the Morgan’s P test, which is typically employed for examining Irish soil P status (Daly & Casey, 2005). 5 mL of dried soil were mixed with 25 mL Morgan’s extractant solution (pH 4.8) (Peech and English 1944) on a horizontal shaker (GFL 3018, Burgwedel, Germany) for 30 min at 180 rpm. Then the solution was filtered using Grade 113 Cellulose qualitative filter paper (Fisherbrand, Fisher Scientific, UK). 1 mL of eluate was incubated with 4 mL colour development solution (Murphy & Riley, 1962) for 30 min, then the absorption of the antimony-phospho-molybdate complex was measured at 882 nm using a spectrophotometer (M501 Single Beam Scanning UV/Visible Spectrophotometer, Camspec, UK). The orthophosphate concentration was calculated via a standard curve prepared with Morgan’s extractant and a 50 mg P L^-1^ potassium dihydrogen phosphate solution to obtain standards with a concentration of 0 (blank), 0.2, 0.5, 1, 2, 3, 4, 5, 6, 7 and 8 mg P L^-1^. The results were compared to the soil P index for grasslands (Plunkett & Wall, 2016).

DNA extraction from 0.25 g rhizosphere soil (frozen at −80°C) was performed using the DNeasy PowerSoil Pro kit (QIAGEN GmbH, Hilden, Germany) according to manual. Quantification of DNA extracts was carried out using the Qubit Fluorometer (Life Technologies, Carlsbad, CA, USA) with a Qubit dsDNA HS assay kit (Life Technologies, Carlsbad, CA, USA). 16S rRNA PCR for denaturing gradient gel electrophoresis (DGGE) was carried out with the primer pair 341F-GC (5‘-CGC CCG CCG CGC GCG GCG GGC GGG GCG GGG GCA CGG GGG GCC TAC GGG AGG CAG CAG-3‘) and 518R (5‘-ATT ACC GCG GCT GCT GG-3‘) developed by Muyzer, de Waal, and Uitterlinden (1993), which targets a 233 bp long gene fragment of the V3 region of the bacterial 16S rDNA gene. The addition of a GC-clamp to the forward primer improves stability in fragment separation during DGGE. In detail, a 25 µL reaction contained 1x DreamTaq buffer (2 mM MgCl_2_, Fisher Scientific, Waltham, MA, USA), 1 M betaine, 2 mM dNTP mix, 0.4 mM of each primer, 0.5 U DreamTaq polymerase (Fisher Scientific, Waltham, MA, USA) and 0.5 µL DNA template. The PCR touchdown protocol was applied in a Mini Amp Plus thermal cycler (Applied Biosystems, Thermo Fisher Scientific, Waltham, MA, USA) as follows: an initial denaturation at 95°C for 5 min, 20 cycles of touchdown annealing from 65 – 55°C (temperature lowered by 0.5°C every cycle), followed by 18 cycles of denaturation at 94°C for 45 s, annealing at 55°C for 45 s and elongation at 72°C for 60 s. Final extension was performed at 72°C for 10 min (Schmalenberger *et al*. 2013). A DGGE fingerprinting analysis was performed as described previously in Fox *et al*. (2014) in a 10 % v/v polyacrylamide gel prepared in 1x TAE buffer in TV-400 DGGE system (Scie-plas, Cambridge, UK), with urea/formamide denaturing gradient of 35-65 % for the 16S rRNA analysis (for preparation details see chapter **Error! Reference source not found.** in the appendix). Band patterns were stained with diluted SYBR^TM^ Gold Nucleic Acid Stain (Invitrogen, Thermo Scientific, Leicestershire, UK) and imaged using a transilluminator (G:box, Syngene).

Amplification of the V4 region of the 16S rRNA gene was carried out in triplicates with primers containing illumina adapters (515F-Illumina 5’-TCG TCG GCA GCG TCA GAT GTG TAT AAG AGA CAG GTG CCA GCM GCC GCG GTA A-3’ and 806R-Illumina 5’-GTC TCG TGG GCT CGG AGA TGT GTA TAA GAG ACA GGG ACT ACH VGG GTW TCT AAT-3’). Each 25 µL reaction contained 1x KAPA HiFi HotStart ReadyMix (2.5 mM MgCl_2_, Roche, CA, USA), 0.3 µM of each primer and 0.5 µL DNA template and the PCR conditions were as follows: initial preincubation at 95°C for 3 min, followed by 25 cycles of denaturation at 98°C for 20 s, annealing at 65°C for 15 s and elongation at 72°C for 15 s, with a final extension step at 72°C for 60 s. The PCR products were purified using the GenElute PCR Clean-up kit (Sigma Aldrich, St. Louis, MO, USA) according to instruction manual, eluted in 50 µL and then 20 µL were sent to CIBIO-InBIO GenomePortugal Research Infrastructure (Vairão, Portugal), where libraries were prepared. Illumina paired[end next-generation sequencing (NGS) was then performed using a 500-cycle Rapid Run kit (Illumina) on a Hiseq2500 sequencer operated by Genewiz (Leipzig, Germany). Demultiplexing was also performed by the sequencing provider.

Sequencing of the functional *phoD* gene fragment was carried out at the University of Minnesota Genomics Centre (UMGC, MN, USA) as PE300 MiSeq amplicon sequencing run. First, the *phoD* gene fragment was amplified with the *phoD* primers *phoD*-F733 (5’-TGG GAY GAT CAY GAR GT-3’) and *phoD*-R1083 (5’-CTG SGC SAK SAC RTT CCA-3’) (Ragot, Kertesz and Bünemann 2015), which provide a higher *phoD* coverage and diversity compared to similar primers published for alkaline phosphatase amplification earlier by Sakurai et al. (2008). The KAPA2G Robust HotStart PCR kit (KAPA Biosystems, Roche) was used and each 25 µL reaction contained 1x KAPA2G Buffer A, 1 M betaine, 0.5 mM MgCl_2_, 0.2 mM dNTPs, 0.8 mM of each primer 0.5 U KAPA2G Robust HotStart DNA Polymerase and 0.5 µL DNA template. The reaction conditions were as follows: 3 min of initial denaturation at 95°C, followed by 35 cycles of denaturation at 95°C for 30 s, annealing at 58°C for 15 s, elongation at 72°C for 15 s and a final extension step at 72°C for 3 min. PCR products were purified using the GenElute PCR Clean-up kit (Sigma Aldrich, St. Louis, MO, USA) according to instruction manual, the triplicates were pooled and diluted 1:5 with PCR water and then subjected to a second PCR adding the illumina adapter sequences. In this PCR the KAPA HiFi HotStart Readymix (KAPA Biosystems, Roche) was used to amplify the samples in triplicates. Each 25 µL reaction contained 1x KAPA HiFi HotStart Readymix, 0.5 mM MgCl_2_, 0.4 mM of each primer and 1 µL of each template. The reaction conditions were 3 min initial denaturation at 95°C, followed by 15 cycles of denaturation at 95°C for 20 s, annealing at 65°C for 15 s and elongation at 72°C for 15 s, and a final extension at 72°C for 1 min. The PCR products were pooled, purified with the GenElute PCR Clean-up kit (Sigma Aldrich) again, and quantified using a Qubit Fluorometer (Life Technologies, Carlsbad, CA, USA) with a Qubit dsDNA HS assay kit (Life Technologies, Carlsbad, CA, USA), before sending the samples to UMGC for library preparation and sequencing and subsequent demultiplexing of sequences.

*phoC* qPCR was conducted with the primers *phoC*-A-F1 (5‘-CGG CTC CTA TCC GTC CGG-3‘) and *phoC*-A-R1 (5’-CAA CAT CGC TTT GCC AGT G-3’) reported in Fraser et al. (2017) on a Roche LightCycler^®^ 96 (Roche Diagnostics, Mannheim, Germany) using the KAPA SYBR FAST qPCR Master Mix (KAPA Biosystems, Cape Town, South Africa). Each 10 µL reaction contained 1x KAPA SYBR FAST qPCR Master Mix (2.5 mM MgCl_2_), 0.3 mM of each primer and 1 µL of DNA template (diluted to 20 ng µL^-1^). The qPCR program started with an initial preincubation at 95 °C for 5 min, then 45 cycles of denaturation at 95 °C for 3 s, primer annealing at 60 °C for 20 s and elongation at 72 °C for 10 s followed. *phoD* qPCR was carried out using the ALPS primers ALPS-F730 (5’-CAG TGG GAC GAC CAC GAG GT-3’) and ALPS-R1101 (5’-GAG GCC GAT CGG CAT GTC G-3’) developed by Sakurai et al. (2008), also applying the KAPA SYBR FAST qPCR Master Mix (KAPA Biosystems) on the LightCycler^®^ 96 (Roche). Each 10 µL reaction contained 1x KAPA SYBR FAST qPCR Master Mix (2.5 mM MgCl_2_), 0.3 mM of each primer and 1 µL of DNA template (diluted to 20 ng µL^-1^). The qPCR program was set up as follows: a 5 min preincubation at 95°C, followed by 45 cycles of denaturation at 95°C for 3 s, annealing at 60°C for 20 s and extension and data acquisition at 72°C for 20 s (adapted from Fraser et al., 2015). The values obtained for *phoC* and *phoD* qPCR were then transformed into copies g^-1^ soil in order to normalize for the DNA extraction yield.

### Statistical Analyses

The results obtained from the shoot dry weight, pH measurements, MPN and CFU analysis, soil ACP and ALP enzymatic assay, Morgan’s P test, *phoD* and *phoC* gene fragment quantification were analysed for statistical significance with the SPSS software (SPSS Statistics, IBM, Version 26). Because the number of samples was lower than 50, normality was checked using Shapiro-Wilk at P>0.05. The homogeneity of variance was evaluated using Levene’s test at P>0.05 and if both assumptions were fulfilled, a one-way analysis of variance (ANOVA) was performed, applying the Tukey HSD post-hoc test for pairwise comparison of the treatments (P<0.05) to assess significant differences between treatment means. Data violating one or both assumptions of normality and homogeneity of variance were transformed either using logarithm or square root transformation, then the ANOVA was repeated, and the untransformed results were reported. When Levene’s Test gave results with P<0.05, the Games-Howell test was applied instead of the Tukey post-hoc analysis. If normality of the data was not achieved, a non-parametric Kruskal-Wallis test was performed instead.

DGGE gel images were compared in Phoretix 1D software (Totallab, Newcastle, UK), creating unweighted pair group method with arithmetic mean (UPGMA) dendrograms. The resulting binary matrix was used in a canonical correspondence analysis (CCA), testing effects of measured environmental factors and visualizing the CCA biplot using the R packages vegan, mabund, permute and lattice, applying a permutational multivariate analysis of variance (PERMANOVA) test using 999 permutations (R Studio, Version 4.0.3).

Demultiplexed, paired-end 16S rRNA sequence reads obtained from CIBIO-InBIO were imported into QIIME2 2020.8(Bolyen *et al*., 2019). The paired-end reads were joined, quality filtered and denoised via the q2-dada2 plugin (Callahan *et al*., 2016). The resulting amplicon sequence variants (ASV) were aligned using the q2-alignment with mafft (Katoh & Standley, 2013) and a phylogenetic tree was built via q2-phylogeny with fasttree2 (Price *et al*., 2010). Samples were rarefied to 14,211 sequences per sample and the alpha diversity metrics (observed species and Faith’s Phylogenetic diversity (Faith, 1992) and beta diversity metrics (Bray-Curtis dissimilarity, Jaccard distance, unweighted UniFrac (Lozupone & Knight, 2005) and weighted UniFrac (Lozupone *et al*., 2007)), and Principle Coordinate Analysis (PCoA) were assessed using the q2-diversity plugin. Taxonomy assignment was accomplished using the q2-feature-classifier (Bokulich *et al*., 2018), training with a SILVA 13.8 99% reference data set for 16S rRNA (Quast *et al*., 2013, Yilmaz *et al*., 2014). First, the reads in the reference data set were trimmed to the region of interest using the primers that had been used in PCR amplification prior to the sequencing. Then, a Naïve Bayes classifier was trained with the trimmed reference sequences and their reference taxonomies. The mvabund and vegan packages in the R software (Version 4.0.3) were used to perform a CCA analysis based on the ASV table obtained during the QIIME2 analysis and the results in the ASV table were visualized as described above for the DGGE analysis. Additionally, taxa bar plots were created to disclose the percentage relative abundance distribution of prevalent taxa among the treatments. Differential abundance was tested using Kruskal-Wallis rank sum test with the function kruskal.test in R Studio (Version 4.0.3) and Wilcoxon post-hoc analysis for multiple pairwise comparisons between groups (function pairwise.wilcox.test), applying Benjamini-Hochberg correction for multiple testing.

Raw *phoD* sequences obtained using the illumina platform were made available demultiplexed by the University of Minnesota Genomics Center. Primers and poor-quality sequences were removed using cutadapt software (Martin, 2011). Usearch (Edgar, 2010) was used to merge paired end reads of the PE300 *phoD* amplicon data. Then the reads were filtered via the fastq_filter command, unique sequences were selected, and duplicates removed via the fastx_uniques command and the UPARSE pipeline was used to cluster sequences into centroid operational taxonomic units (OTUs) (Edgar, 2013). A 75% sequence similarity threshold was used for OTU clustering (Fraser *et al*., 2015). A reference database for the alkaline phosphomonoesterase gene *phoD* was downloaded from the fungene functional gene repository (Fish *et al*., 2013) and was used as database to assign taxonomy. The OTU table was not rarefied to avoid omission of rare OTUs and decreased sensitivity due to an increase in Type II errors (McMurdie & Holmes, 2014). The mvabund and vegan packages in the R software (Version 4.0.3) were used to create a CCA or non-metric-multidimensional scaling (NMDS) biplot and subsequent permutation based on the OTU table obtained during the Usearch analysis and the results in the ASV table were visualized as described above for the DGGE analysis. Additionally, taxa bar plots were created to disclose the percentage relative abundance distribution of prevalent genera among the treatments, as described above for the 16S NGS data.

## Results

### Shoot Biomass Yield, Agronomic Efficiency and Elemental Analysis of Ryegrass Biomass

Fresh biomass records showed a slight trend for higher yields in treatments with a higher P application rate (see Table 3). Dry biomass yield of the *L. perenne* shoots was low for all treatments, ranging between 1.9 and 3.7 g per pot. The average dry weight values of the high P fertilized corrected for the seed germination rate (dry weight to shoot count ratio) treatments were higher than the ones with low P fertilization, with one exception: MWS20 yielded a higher dry weight per shoot count ratio than MWS60. The PLA20 treatment displayed the lowest values of all treatments, even performing poorer that the P-free control SP0. Generally, both struvites yielded the highest plant dry weight per shoot count on average, with MWS20 and CGS60 being the only treatments with significantly higher (P<0.05) yields than the P-free control. The P utilization efficiency was evaluated as agronomic efficiency (P-uptake/P-supplied; AE), because due to the low dry matter yield other means of evaluation were not possible. Because of its low dry matter yield, averaging below SP0, the AE for PLA20 is negative. Interestingly, the low P input treatments SP20, SSA20 and MWS20 resulted in a higher AE that their high P input equivalents in the short term.

**Table 3.**
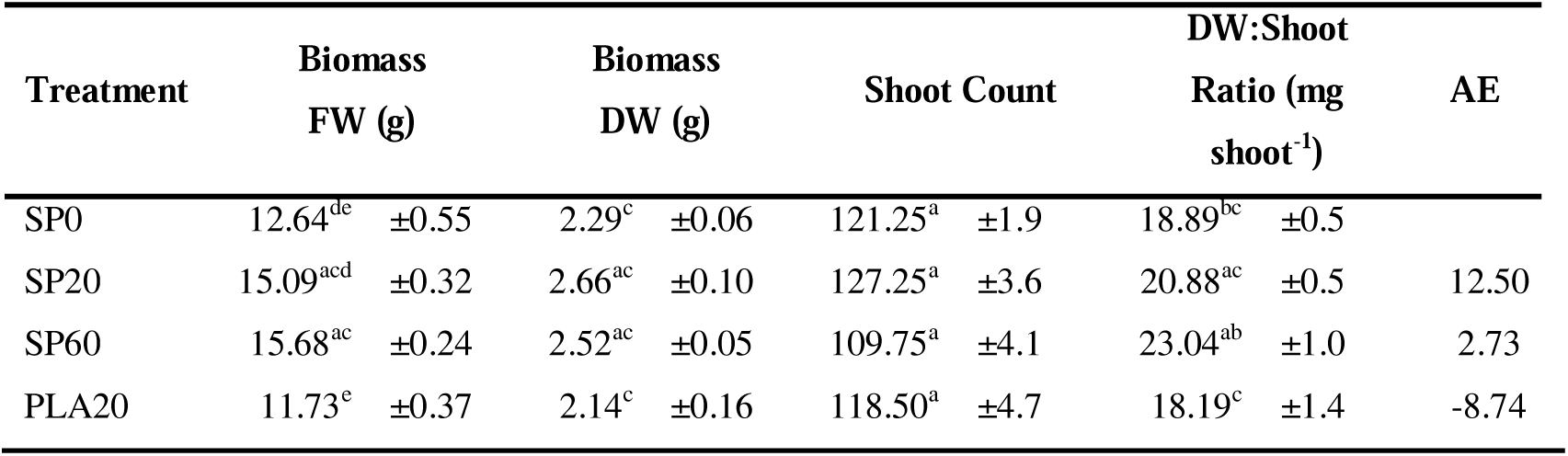
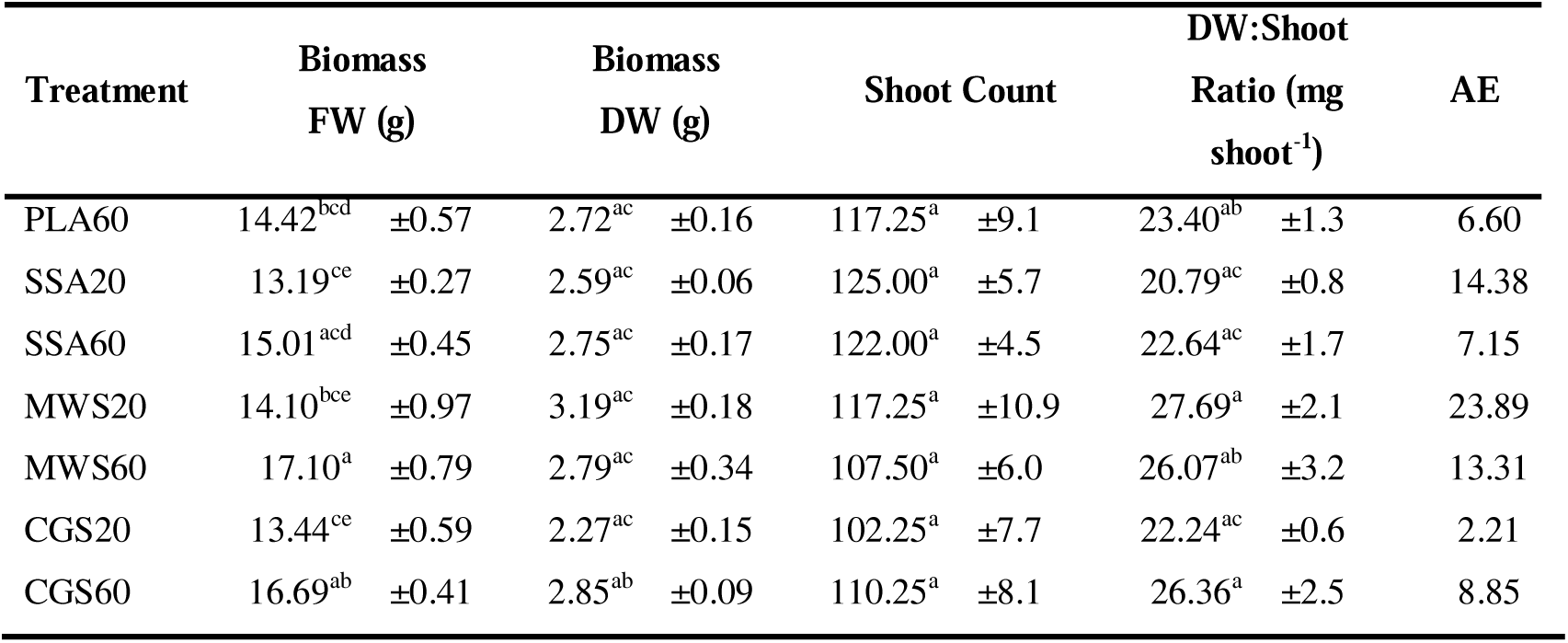
Average values for fresh weight (FW) and dry weight (DW) of Lolium perenne harvest, shoot count, dry weight to shoot ratio, measured in quadruplicates after 54 days of growth in pots, and agronomic efficiency (AE), different letters indicate significant difference (P < 0.05) within a column, determined via one-way ANOVA with Tukey HSD or Games-Howell post-hoc analysis, ± represents standard error.

The mass balance of the macro-(Table 4) and micronutrients (Table 5) could only be determined for the pooled treatments. Therefore, because only one measurement was obtained per treatment, statistical significance between the treatments could not be evaluated. Interestingly, Ca, K and N uptake was highest in PLA20, which performed lowest in terms of dry biomass yield, and a similar pattern is visible for CGS20, which also exhibited low grass yields. Mg and S were similar between all treatment. P uptake was highest in SP60, and all treatment except for PLA show higher P uptake in the high P fertilized treatments compared to the low P application rate of the same fertilizer.

**Table 4.** Mass balance of total perennial ryegrass shoot dry weight (pooled per treatment), in µg per treatment (P added as 0, 20 or 60 kg ha^-1^), for the macronutrients of the pot trial after 54 days of growth in a greenhouse.

| Treatment | Macronutrients (µg treatment<sup>-1</sup>) |  |  |  |  |  |
| --- | --- | --- | --- | --- | --- | --- |
|  | Calcium | Magnesium | Sulphur | Nitrogen | Phosphorus | Potassium |
| SP0 | 1619.8 | 371.2 | 360.0 | 4713.2 | 258.7 | 4836.9 |
| SP20 | 1397.2 | 329.3 | 339.3 | 4760.5 | 299.4 | 4471.1 |
| SP60 | 1360.3 | 321.9 | 363.4 | 5077.9 | 415.4 | 4911.7 |
| PLA20 | 1777.8 | 407.4 | 419.8 | 5530.9 | 296.3 | 5370.4 |
| PLA60 | 1245.3 | 290.3 | 318.4 | 4307.1 | 252.8 | 4194.8 |
| SSA20 | 1275.8 | 294.4 | 323.8 | 4367.0 | 235.5 | 4210.0 |
| SSA60 | 1135.7 | 267.8 | 313.9 | 4478.3 | 295.5 | 3942.8 |
| MWS20 | 1194.6 | 280.5 | 298.6 | 4461.5 | 262.4 | 4181.0 |
| MWS60 | 944.0 | 272.0 | 264.0 | 3944.0 | 304.0 | 3576.0 |
| CGS20 | 1507.2 | 363.0 | 374.0 | 5181.5 | 308.0 | 5005.5 |
| CGS60 | 992.0 | 301.2 | 292.3 | 4242.7 | 336.6 | 3950.4 |

**Table 5.** Mass balance of total perennial ryegrass shoot dry weight (pooled per treatment), in µg per treatment(P added as 0, 20 or 60 kg ha^-1^), for the micronutrients of the pot trial after 54 days of growth in a greenhouse.

| Treatment | Micronutrients |  |  |  |  |  |
| --- | --- | --- | --- | --- | --- | --- |
|  | Boron | Copper | Iron | Manganese | Molybdenum | Zinc |
| SP0 | 0.9 | 5.5 | 217.8 | 36.2 | 0.01 | 6.5 |
| SP20 | 0.6 | 3.3 | 116.8 | 30.7 | 0.02 | 5.8 |
| SP60 | 0.6 | 3.1 | 151.7 | 27.7 | 0.02 | 6.2 |
| PLA20 | 1.0 | 6.1 | 234.0 | 32.8 | 0.02 | 6.6 |
| PLA60 | 0.6 | 2.7 | 146.3 | 18.5 | 0.01 | 5.3 |
| SSA20 | 0.7 | 3.5 | 193.8 | 23.9 | 0.01 | 5.1 |
| SSA60 | 0.6 | 3.0 | 139.8 | 17.1 | 0.01 | 5.4 |
| MWS20 | 0.6 | 2.9 | 144.1 | 24.1 | 0.01 | 4.3 |
| MWS60 | 0.4 | 2.3 | 102.0 | 21.0 | 0.01 | 3.8 |
| CGS20 | 0.7 | 2.9 | 158.0 | 30.6 | 0.01 | 5.5 |
| CGS60 | 0.5 | 2.9 | 122.0 | 23.6 | 0.01 | 4.8 |

The most abundant micronutrient was iron; its uptake was prominently higher in SP0 and PLA20, the treatments with the lowest dry biomass yield. Fe was followed by Mn, Zn and C in abundance, B and Cu were detected at particularly low concentrations.

### RDF Influence on P Mineralising and P Solubilizing Microbial Community

The MPN approach yielded no statistically significant difference in P mobilization capabilities from phosphonoacetic acid and phytate and overall heterotrophic bacteria, as shown in Table 6. Nonetheless, average MPN g^-1^ soil of PLA60, SSA60 and CGS60 were around twice as high or higher than for SP60 in terms of PAA mobilization. For the TCP solubilization a trend was visible, indicating that mineral P fertilizer had a lower CFU count for TCP solubilizing colonies compared to the RDFs. The SP60 treatment had the lowest CFU counts per gram soil and was also significantly lower (P<0.05) than the RDFs PLA60, MWS20 and MWS60.

**Table 6.** MPN values of phosphonoacetic acid mobilizing (PAA)and phytate mobilizing (Phy) bacteria, and MPN values of total heterotrophic bacteria (R2A) and CFU values of tri-calcium phosphate solubilizing bacteria (TCP), different letters indicate significant difference (P < 0.05) within a column, determined via one-way ANOVA with Tukey HSD or Games-Howell post-hoc analysis, ± represents standard error, n=4.

| Treatment | PAA<br>(MPN g<sup>-1</sup> soil) |  | Phy<br>(MPN g<sup>-1</sup> soil) |  | R2A<br>(MPN g<sup>-1</sup> soil) |  | TCP<br>(CFU g<sup>-1</sup> soil) |  |
| --- | --- | --- | --- | --- | --- | --- | --- | --- |
| SP0 | 3.1E+06<sup>a</sup> | $\pm 1.0E+06$ | 18.6E+06<sup>a</sup> | $\pm 5.6E+06$ | 92.5E+06<sup>a</sup> | $\pm 2.5E+07$ | 275.0E+03<sup>bcd</sup> | $\pm 3.1E+04$ |
| SP20 | 7.5E+06<sup>a</sup> | $\pm 3.0E+06$ | 11.5E+06<sup>a</sup> | $\pm 3.5E+06$ | 141.3E+06<sup>a</sup> | $\pm 3.5E+07$ | 150.0E+03<sup>cd</sup> | $\pm 3.2E+04$ |
| SP60 | 6.7E+06<sup>a</sup> | $\pm 1.2E+05$ | 8.3E+06<sup>a</sup> | $\pm 2.9E+06$ | 120.0E+06<sup>a</sup> | $\pm 1.9E+07$ | 138.0E+03<sup>d</sup> | $\pm 3.6E+04$ |
| PLA20 | 9.8E+06<sup>a</sup> | $\pm 2.6E+06$ | 19.8E+06<sup>a</sup> | $\pm 7.0E+06$ | 218.8E+06<sup>a</sup> | $\pm 5.1E+07$ | 250.0E+03<sup>bcd</sup> | $\pm 5.0E+04$ |
| PLA60 | 14.2E+06<sup>a</sup> | $\pm 5.9E+06$ | 11.3E+06<sup>a</sup> | $\pm 1.2E+05$ | 102.5E+06<sup>a</sup> | $\pm 2.4E+07$ | 1.0E+06<sup>a</sup> | $\pm 7.6E+05$ |
| SSA20 | 5.2E+06<sup>a</sup> | $\pm 7.4E+05$ | 31.3E+06<sup>a</sup> | $\pm 1.3E+07$ | 154.5E+06<sup>a</sup> | $\pm 4.9E+07$ | 850.0E+03<sup>ad</sup> | $\pm 1.6E+05$ |
| SSA60 | 10.1E+06<sup>a</sup> | $\pm 4.4E+06$ | 12.8E+06<sup>a</sup> | $\pm 9.4E+05$ | 156.3E+06<sup>a</sup> | $\pm 3.4E+07$ | 488.0E+03<sup>ad</sup> | $\pm 4.4E+05$ |
| MWS20 | 5.4E+06<sup>a</sup> | $\pm 6.5E+05$ | 19.3E+06<sup>a</sup> | $\pm 3.2E+06$ | 156.3E+06<sup>a</sup> | $\pm 2.7E+07$ | 925.0E+03<sup>ab</sup> | $\pm 8.4E+04$ |
| MWS60 | 9.5E+06<sup>a</sup> | $\pm 4.5E+06$ | 16.0E+06<sup>a</sup> | $\pm 2.0E+05$ | 132.5E+06<sup>a</sup> | $\pm 2.0E+07$ | 863.0E+03<sup>ac</sup> | $\pm 9.7E+04$ |
| CGS20 | 6.2E+06<sup>a</sup> | $\pm 2.4E+05$ | 19.8E+06<sup>a</sup> | $\pm 6.7E+06$ | 102.5E+06<sup>a</sup> | $\pm 2.4E+07$ | 575.0E+03<sup>ad</sup> | $\pm 9.4E+04$ |
| CGS60 | 12.0E+06<sup>a</sup> | $\pm 1.7E+06$ | 15.3E+06<sup>a</sup> | $\pm 1.3E+06$ | 61.7E+06<sup>a</sup> | $\pm 2.4E+06$ | 763.0E+03<sup>ad</sup> | $\pm 8.7E+04$ |

The pH-CaCl_2_ was significantly higher for all fertilized treatments compared to SP0, as shown in Table 7. It is noteworthy that the pH value was higher for the higher P application rate of each fertilizer compared to the lower dose. The ash fertilizers PLA and SSA had a significantly higher liming effect on the soil compared to the other treatments at their high P application rate, followed by the mineral SP fertilizer and finally the struvite RDFs, which were affecting the soil pH the least. The Morgan’s P test on the other hand revealed highest P availability in the struvite treatments, however, due to high variability of the results and small replicate number, there was no significant difference to other fertilized treatments. However, while the ash RDFs and the mineral fertilizers averaged in an index 1 range, the treatments MWS60 and CGS60 averaged at index 2 and index 3 range concentrations, respectively. The high variability of the Morgan’s P results could stem from the RDF properties. While the ashes consisted of a fine powdery structure, the struvites were received as granules and therefore they were not as well distributed throughout the soil. This could have led to some soil samples containing more granules than other replicates, causing a higher variability of the results due to higher P fertilizer input.

**Table 7.** Mean values for soil pH and bioavailable P measured using Morgan’s P test after 54 days of growth, different letters indicate significant difference (P < 0.05) within a column, determined via one-way ANOVA with Tukey HSD or Games-Howell post-hoc analysis, ± represents standard error, n=4.

| Treatment | Soil pH | Morgan's P<br>(mg P L<sup>-1</sup>) |
| --- | --- | --- |
| SP0 | 4.75<sup>d</sup> $\pm 0.03$ | 0.58<sup>c</sup> $\pm 0.08$ |
| SP20 | 4.94<sup>bc</sup> $\pm 0.02$ | 1.01<sup>ac</sup> $\pm 0.12$ |
| SP60 | 5.05<sup>b</sup> $\pm 0.04$ | 2.35<sup>ab</sup> $\pm 0.26$ |
| PLA20 | 4.95<sup>bc</sup> $\pm 0.02$ | 1.98<sup>ab</sup> $\pm 0.17$ |
| PLA60 | 5.22<sup>a</sup> $\pm 0.03$ | 2.75<sup>b</sup> $\pm 0.19$ |
| SSA20 | 4.96<sup>bc</sup> $\pm 0.02$ | 1.57<sup>ab</sup> $\pm 0.11$ |
| SSA60 | 5.24<sup>a</sup> $\pm 0.02$ | 2.48<sup>ac</sup> $\pm 0.41$ |
| MWS20 | 4.87<sup>cd</sup> $\pm 0.04$ | 2.00<sup>abc</sup> $\pm 0.33$ |
| MWS60 | 5.02<sup>b</sup> $\pm 0.02$ | 4.46<sup>abc</sup> $\pm 0.60$ |
| CGS20 | 4.90<sup>c</sup> $\pm 0.02$ | 2.79<sup>abc</sup> $\pm 0.45$ |
| CGS60 | 4.95<sup>bc</sup> $\pm 0.01$ | 5.16<sup>abc</sup> $\pm 0.71$ |

Potential acid phosphomonoesterase (ACP) activity was around 10-fold higher than alkaline phosphomonoesterase (ALP) activity in the rhizosphere soil measured after destructive harvest of the pots, as shown in Table 8. The mineral SP60 fertilizer and the struvite treatments MWS60 and CGS60 demonstrated a positive effect on potential ACP activity, while the ash treatments were not significantly different to the P-free SP0 control, with the exception of SSA20, which had the highest measured activity of the ashes on average. Colorimetric determination of potential ALP activity showed no significant difference between treatments. Table 8 also states the *phoC* and *phoD* gene copy numbers per gram of soil corrected for DNA extraction yields. While there was no significant difference detected in the *phoC* gene copy numbers, a significantly lower (P<0.05) *phoD* gene copy number was found in the PLA20 treatment compared to the CGS20 treatment, which displayed the highest copy number count for *phoD*. Contrasting to the findings of the enzyme assay, the *phoD* gene copy numbers were 1000-fold higher than for *phoC*.

**Table 8.** Average potential acid (ACP) and alkaline phosphomonoesterase (ALP) activity determined via spectrophotometry and phoC and phoD gene copy numbers per gram of soil normalized for DNA extraction yield determined via qPCR after destructive harvest of pot after 54 days of growth, different letters indicate significant difference (P < 0.05) within a column, determined via one-way ANOVA with Tukey HSD or Games-Howell post-hoc analysis, ± represents standard error, n=4.

| Treatment | ACP<br>($\mu\text{g pNP}$<br>$\text{g}^{-1} \text{ soil h}^{-1}$) | ALP<br>($\mu\text{g pNP}$<br>$\text{g}^{-1} \text{ soil h}^{-1}$) | *phoC*<br>(*phoC* copies<br>$\text{g}^{-1} \text{ soil}$) | *phoD*<br>(*phoD* copies<br>$\text{g}^{-1} \text{ soil}$) |
| --- | --- | --- | --- | --- |
| SP0 | 1653.7<sup>d</sup> ± 77.4 | 218.1<sup>a</sup> ± 10.6 | 131.3E+03<sup>a</sup> ± 6.3E+04 | 148.2E+06<sup>ab</sup> ± 5.1E+06 |
| SP20 | 1663.9<sup>d</sup> ± 38.9 | 278.7<sup>a</sup> ± 10.2 | 79.7E+03<sup>a</sup> ± 3.3E+04 | 140.9E+06<sup>ab</sup> ± 2.9E+06 |
| SP60 | 2167.0<sup>abc</sup> ± 78.8 | 259.5<sup>a</sup> ± 16.4 | 98.9E+03<sup>a</sup> ± 3.4E+04 | 140.7E+06<sup>ab</sup> ± 3.1E+06 |
| PLA20 | 2007.1<sup>bd</sup> ± 45.6 | 208.4<sup>a</sup> ± 20.7 | 75.8E+03<sup>a</sup> ± 3.3E+04 | 131.7E+06<sup>b</sup> ± 7.0E+06 |
| PLA60 | 1712.3<sup>cd</sup> ± 81.7 | 217.5<sup>a</sup> ± 7.0 | 119.9E+03<sup>a</sup> ± 5.9E+04 | 151.7E+06<sup>ab</sup> ± 4.0E+06 |
| SSA20 | 2183.4<sup>abc</sup> ± 94.3 | 188.0<sup>a</sup> ± 7.1 | 75.9E+03<sup>a</sup> ± 2.9E+04 | 133.5E+06<sup>ab</sup> ± 6.9E+06 |
| SSA60 | 2084.2<sup>bd</sup> ± 103.5 | 228.8<sup>a</sup> ± 27.2 | 88.3E+03<sup>a</sup> ± 4.5E+04 | 138.4E+06<sup>ab</sup> ± 6.6E+06 |
| MWS20 | 2262.8<sup>bd</sup> ± 137.6 | 233.9<sup>a</sup> ± 33.3 | 100.8E+03<sup>a</sup> ± 3.6E+04 | 145.5E+06<sup>ab</sup> ± 1.7E+07 |
| MWS60 | 2262.8<sup>ab</sup> ± 103.1 | 233.9<sup>a</sup> ± 30.4 | 168.6E+03<sup>a</sup> ± 7.5E+04 | 163.3E+06<sup>ab</sup> ± 5.3E+06 |
| CGS20 | 2066.9<sup>bd</sup> ± 133.7 | 199.9<sup>a</sup> ± 23.2 | 218.4E+03<sup>a</sup> ± 5.0E+04 | 168.9E+06<sup>a</sup> ± 9.8E+06 |
| CGS60 | 2616.4<sup>a</sup> ± 163.7 | 209.0<sup>a</sup> ± 17.9 | 88.2E+03<sup>a</sup> ± 3.3E+04 | 154.3E+06<sup>ab</sup> ± 4.1E+06 |

### DNA Fingerprint Analysis of Bacterial Community Structure and Next Generation Sequencing of Bacterial 16S rRNA Gene

PCR-DGGE was performed for individual treatments PLA20 and PLA60; SSA20 and SSA60; and MWS20, MWS60, CGS20 and CGS60, always in comparison to SP0 and SP60 to analyse the bacterial community structure. UPGMA dendrograms demonstrated high similarity between the band patterns of all treatments, the percentage of similarity was always above 90%, as displayed in 3.

For the 16S rRNA amplicon sequencing, a total of 7,954,388 demultiplexed sequences were received. On average, 180,782 sequences per sample were input into DADA2, 161,072 (86.1%) were filtered, 127,158 sequences per sample (66.3%) were merged and 74,522 (39.4%) of these were detected as non-chimeric. None of the alpha diversity estimators were statistically significantly different. Chao1 values ranged from 864 (PLA60) to 1758 (SP0), while the Shannon index ranged from 5.6 (SSA20) to 7.1 (SP0). The richness estimators ACE, Chao1 and observed features were identical due to the removal of singletons in the ASV matrix output in QIIME2.

The CCA biplot of the 16S rRNA bacterial community analysed via amplicon sequencing displayed limited separation of treatments (Figure 4). Of the treatments, only SP0 was separated on the first axis, while MWS60 was separated on the 2^nd^ axis. Pairwise comparison of treatments using the PERMANOVA as described in the pairwise Adonis function of the vegan package in R revealed no significant differences, when a Benjamini-Hochberg correction was applied. Significant environmental parameters involved in the visible shift of the bacterial soil community compared to the non-P fertilized treatment were soil pH, P plant availability, acid phosphatase activity, TCP solubilizing CFUs and dry weight yields of *L. perenne*.

**Figure 4.**
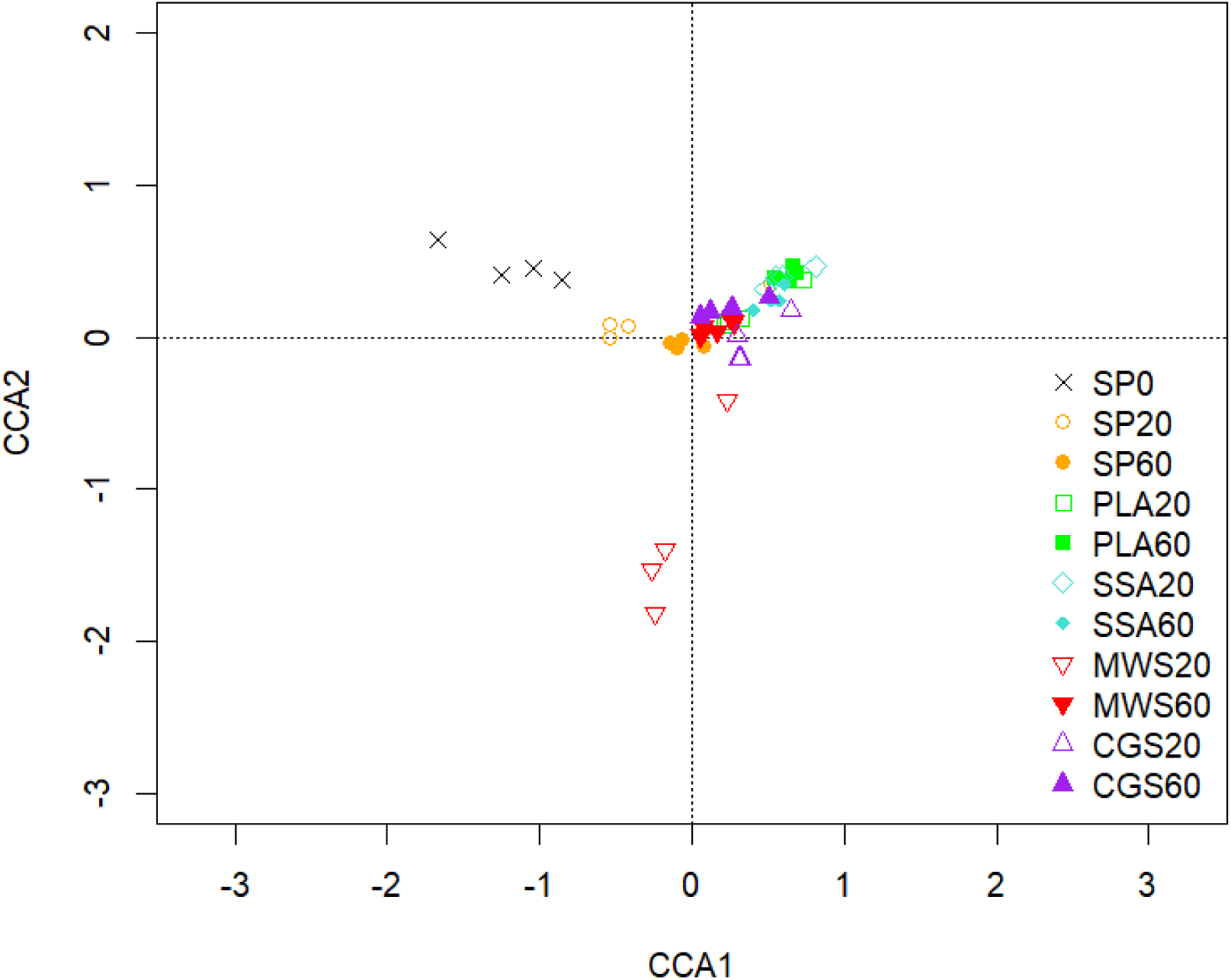
CCA of the bacterial community sequences based on ASVs obtained from 16S rRNA, CCA1 explains 3.44% and CCA2 3.05% of the total variation of the data, n=4.

Figure 5 shows the mean relative abundance of the ten most abundant phyla detected in the 16S rRNA analysis. The phyla *Actinobacteria*, *Planctomycetes*, *Firmicutes*, *Proteobacteria*, *Verrucomicrobia* and *Acidobacteria* comprise over 75% of all phyla.

**Figure 5.**
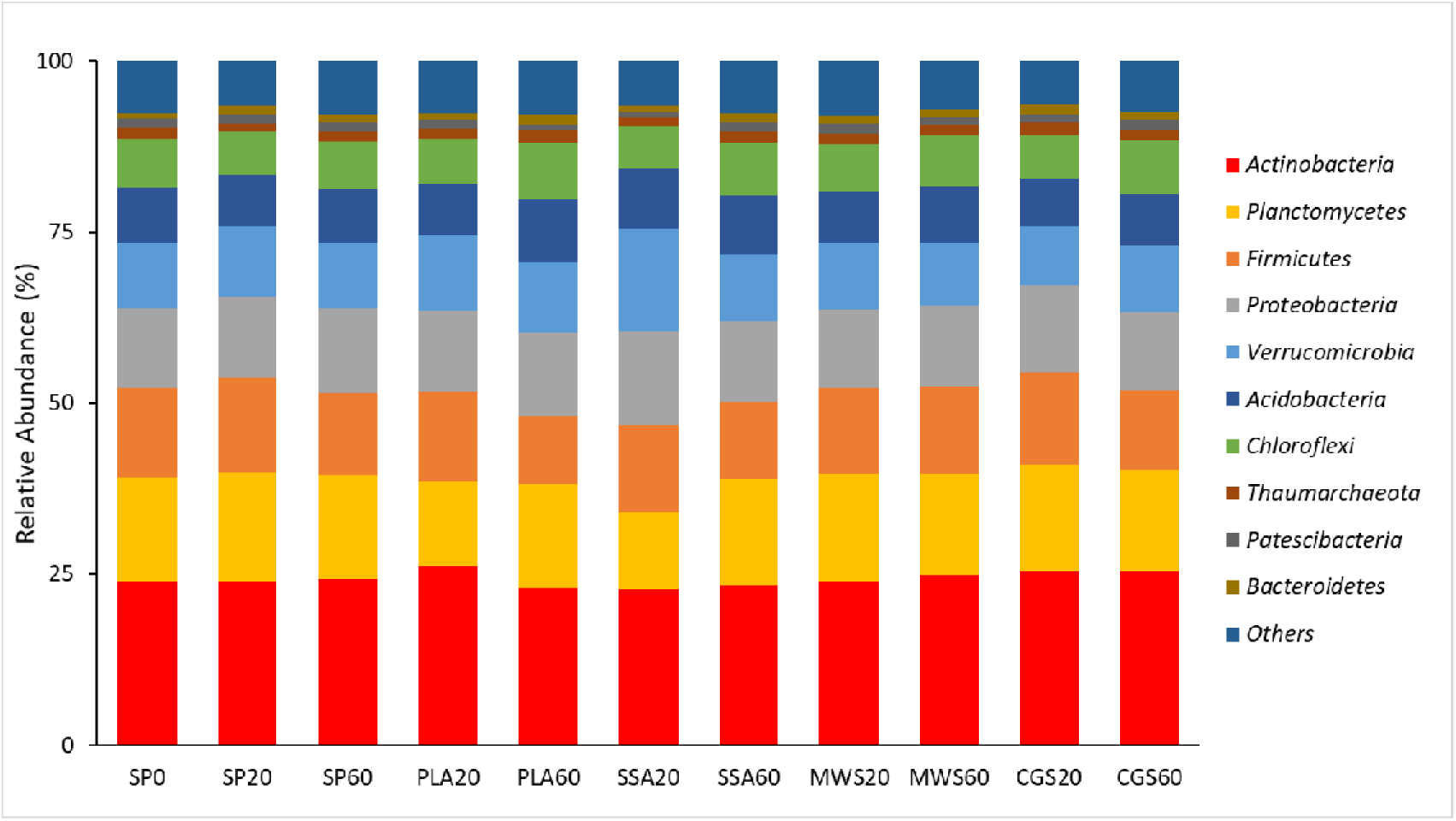
Mean relative abundance of the top 10 abundant phyla of the 16S rRNA amplicon sequencing, n=4.

Out of the 21 phyla of the 16S rRNA sequencing that had a relative abundance across treatments of above 1.5%, five showed statistically significant differences in pairwise post-hoc comparison, as depicted in Figure 6. *Acidobacteria* was significantly higher abundant (P<0.05) in PLA60 in comparison to CGS20. *Patescibacteria* was significantly higher abundant (P<0.05) in the ash treatment SSA60 and the struvite treatments MWS20 and CGS60 compared to PLA60 and SSA20. *Euryarcheota* on the other hand was also significantly higher abundant (P<0.05) in MWS20 than in the SSA ash treatment at both concentrations. The phylum WS4 was significantly higher abundant (P<0.05) in the SSA60 treatment, compared to SP20 and CGS60. *Halanaerobiaeota* had a significantly lower abundance (P<0.05) in all P fertilized treatments compared to the control SP0.

**Figure 6.**
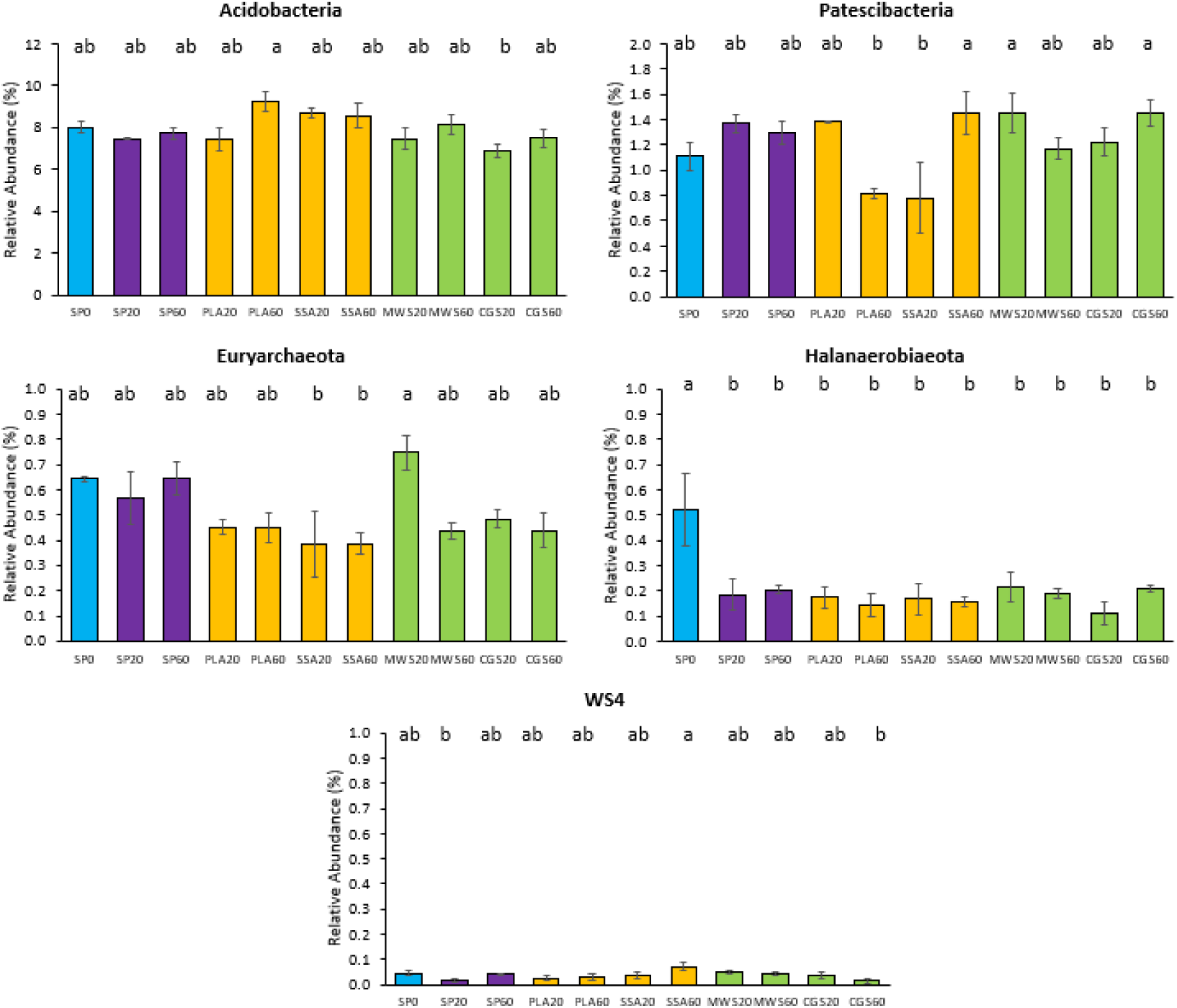
Mean relative abundance of selected phyla of the 16S rRNA sequencing with an abundance above a cut-off of 1.5% which showed significant differences in the treatments, significance determined via Kruskal-Wallis test and Wilcoxon post-hoc analysis with Benjamini-Hochberg correction, different letters indicate significant difference (P < 0.05), n=4.

For the analysis of significant differences within the ASVs assigned down to genus level, genera with an abundance above 20% were selected for Kruskal-Wallis differential abundance analysis. Statistical analysis of all treatments together only revealed significant differences (P<0.05) in the genus *Nocardioides* (SP20 was significantly different to SSA20). The differential abundance analysis was then repeated with subset of treatments, where SP0 was compared to all RDF20, SP0 to all RDF60, SP20 to all RDF20 and SP60 to all RDF60. For the comparisons SP0 vs RDF20 and SP20 vs RDF20, no significant differences were detected in the 20 most abundant genera for the 16S rRNA sequencing analysis. Differential abundance analysis of SP0 vs RDF60 and SP60 vs RDF60 revealed in both cases significant differences (P<0.05) in the genera *Nocardioides*, *Conexibacter*, *Clostridium sensu stricto 13* and an uncultured bacterium belonging to the family *Isosphaeraceae.* The results for the SP0 vs RDF60 and SP60 vs RDF60 Wilcoxon post-hoc analysis were overlapping, with the SP60 vs RDF60 having one additional significant different (P<0.05) treatment pair, therefore only the results for the SP60 vs RDF60 were shown, as summarized in Figure 7 below.

**Figure 7.**
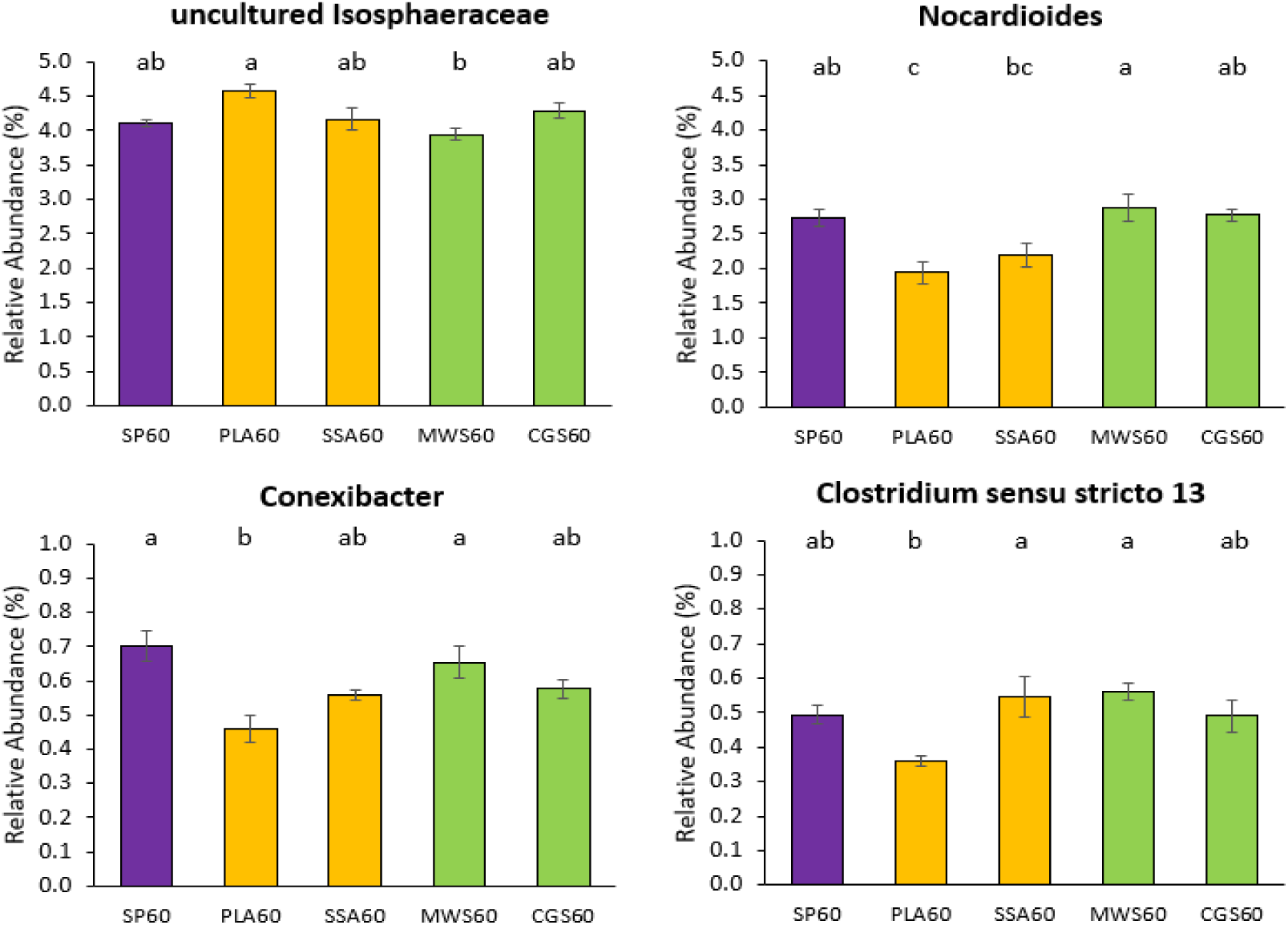
Mean relative abundance of selected genera above the cut-off of 20% overall relative abundance, which were significantly different abundant for the 16S rRNA sequencing analysis, for the separate analysis of the high P treatments SP60, PLA60, SSA60, MWS60 and CGS60 only, significance determined via Kruskal-Wallis test and Wilcoxon post-hoc analysis with Benjamini-Hochberg correction, different letters indicate significant difference, n=4.

The uncultured *Isophaeraceae* had the highest relative abundance in the PLA60 treatment, while the MWS60 demonstrated significantly lower (P<0.05) abundance of this genera compared to PLA60. For the other three genera on the other hand, MWS60 was significantly higher abundant (P<0.05) than most other treatments, while the PLA60 treatment was significantly lower (P<0.05) abundant than all other treatments. For the genus *Conexibacter* also the mineral SP60 treatment showed significantly higher (P<0.05) relative abundance in combination with MWS60, and the genus *Clostridium sensu stricto 13* was significantly higher (P<0.05) abundant in the MWS60 as well as the SSA60 treatment.

### Next-Generation Sequencing Analysis of the phoD Harbouring Bacterial Community in Soil

4 570 420 *phoD* paired-end amplicon sequences were received from the sequencing facility, out of which 2 734 589 (59.8%) were successfully merged using USEARCH. 2 189 710 of the merged sequences (80.1%) were filtered and these sequences contained 1 260 580 (86.3%) singletons. In total, 9 204 OTUs were picked at a 75% similarity threshold and 3 278 chimeral sequences (1.6%) were removed. Finally, 2 356 148 (86.2%) out of 2 734 589 merged sequences were mapped to OTUs.

Alpha diversity analysis via the phyloseq package in R showed no significant difference between the treatments for the estimates observed features, Chao1 (5164-5659), ACE (5291-5878), Shannon (6.4-6.5) and Simpson (0.99). The community structure of *phoD* harbouring bacteria displayed in a CCA biplot (Figure 8) shows a visible separation of the MWS60 treatment from all other treatments on the first axis and a separation of the SSA60 treatment on the second axis. However, PERMANOVA using 999 permutations and subsequent pairwise comparison with Benjamini-Hochberg correction revealed no significant differences (P>0.05) between the treatments.

**Figure 8.**
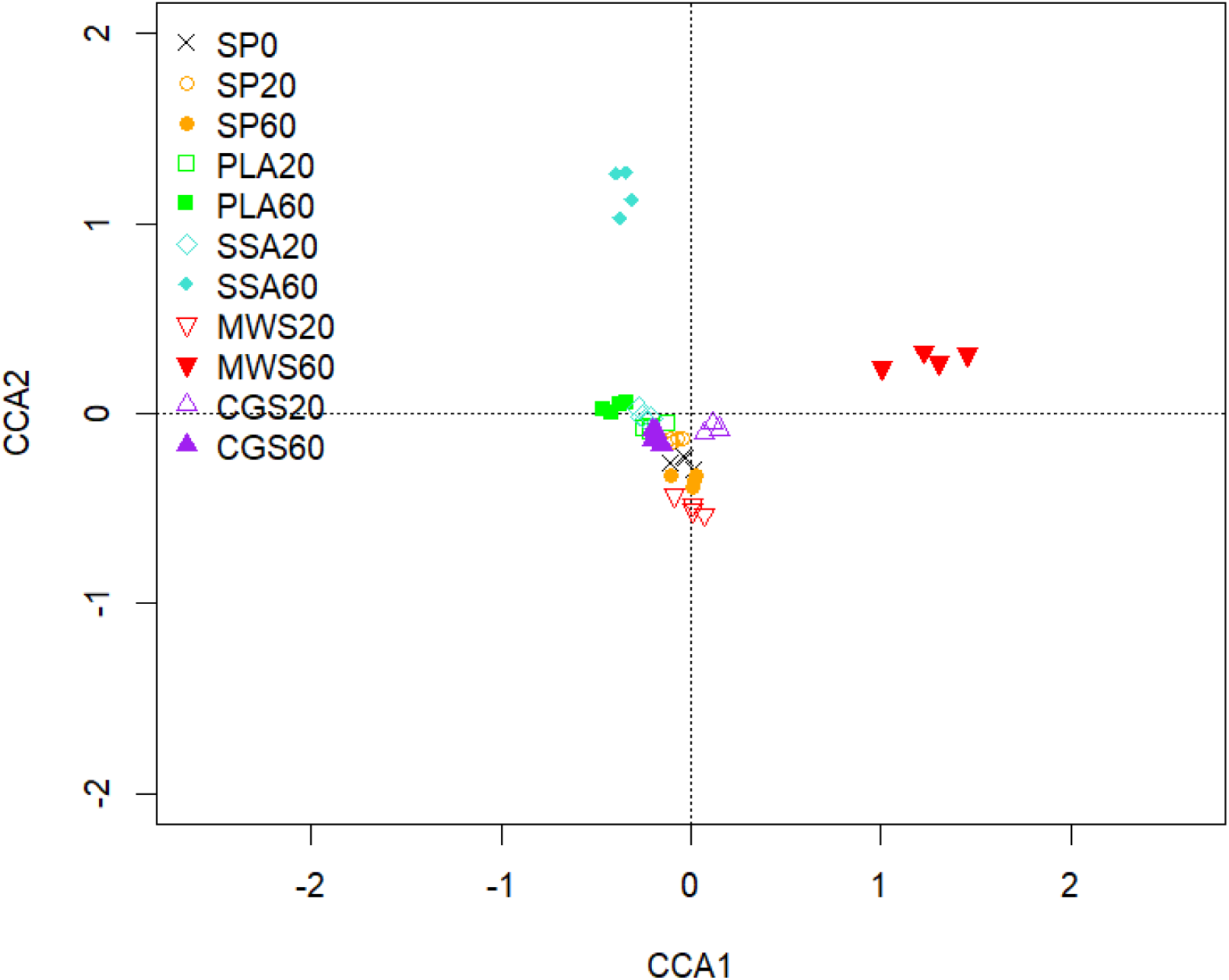
CCA of the phoD gene harbouring community sequences based on OTUs, CCA1 explains 3.04% and CCA2 2.73% of the total variation of the data, n=4.

The ten most abundant genera were Bifidobacterium, Oerskovia, Frigoribacterium, Streptomyces, Xanthomonas, Microbacterium, Rhodopseudomonas, Leifsonia, Rubrivivax and Rhodanobacter, and these genera constituted around 50% of the total genera in the samples (see Figure 9).

**Figure 9.**
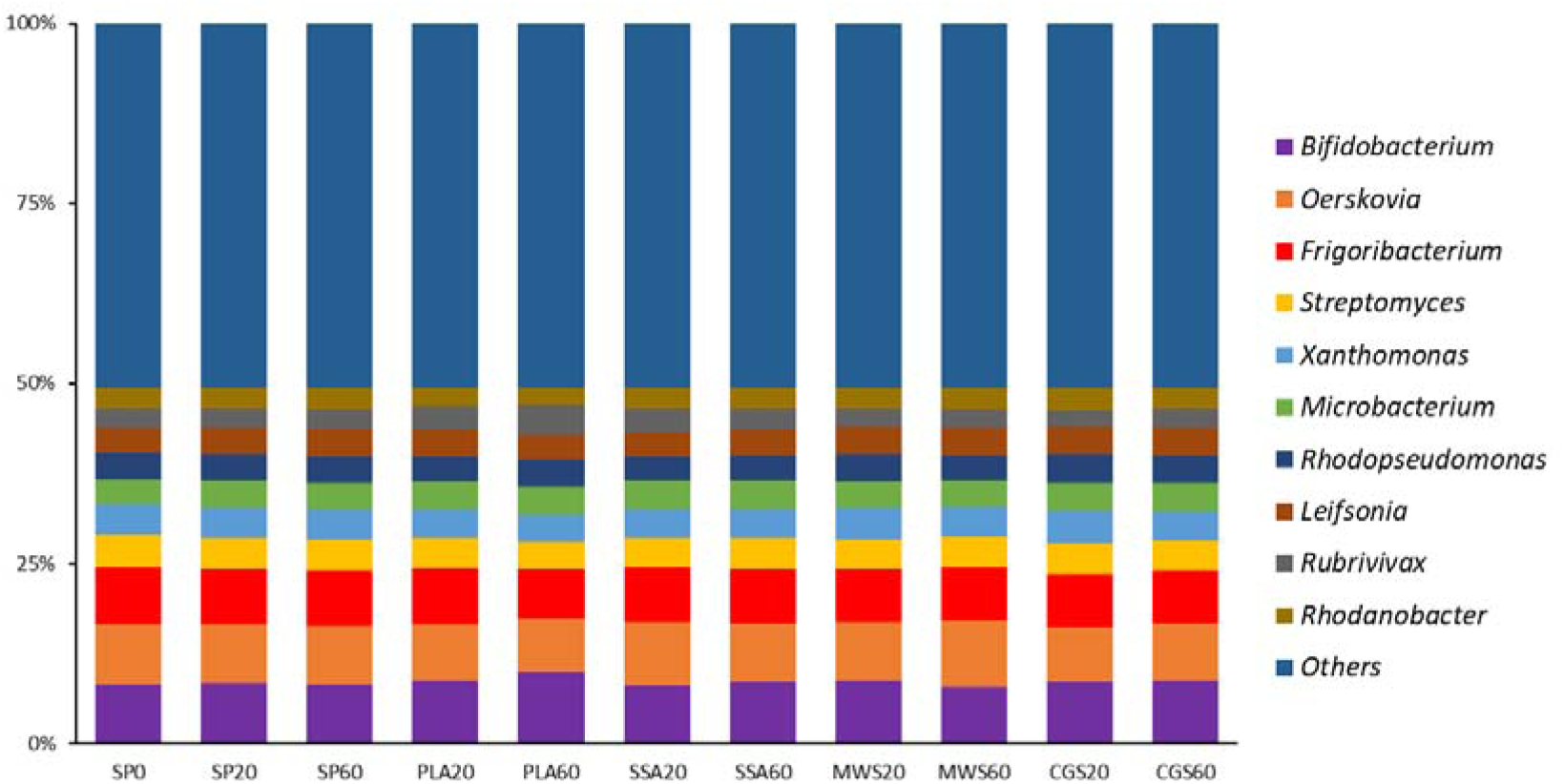
Mean relative abundance of the top 10 abundant genera of the phoD amplicon sequencing of the pot trial, all other genera collapsed in “others”, n=4.

Six of the ten most abundant genera showed significant differences in their abundance in the post-hoc analysis (P<0.05, see Figure 10). The genera *Oerskovia*, *Frigoribacterium*, *Xanthomonas* and *Rhodanobacter* were significantly lower abundant in the PLA60 treatment. *Microbacterium* and *Rubrivivax* on the other hand had a significantly higher (P<0.05) relative abundance in the PLA60 treatment, these genera had generally higher relative abundances in most ash treatments, compared to the SP and struvite treatments MWS and CGS.

**Figure 10.**
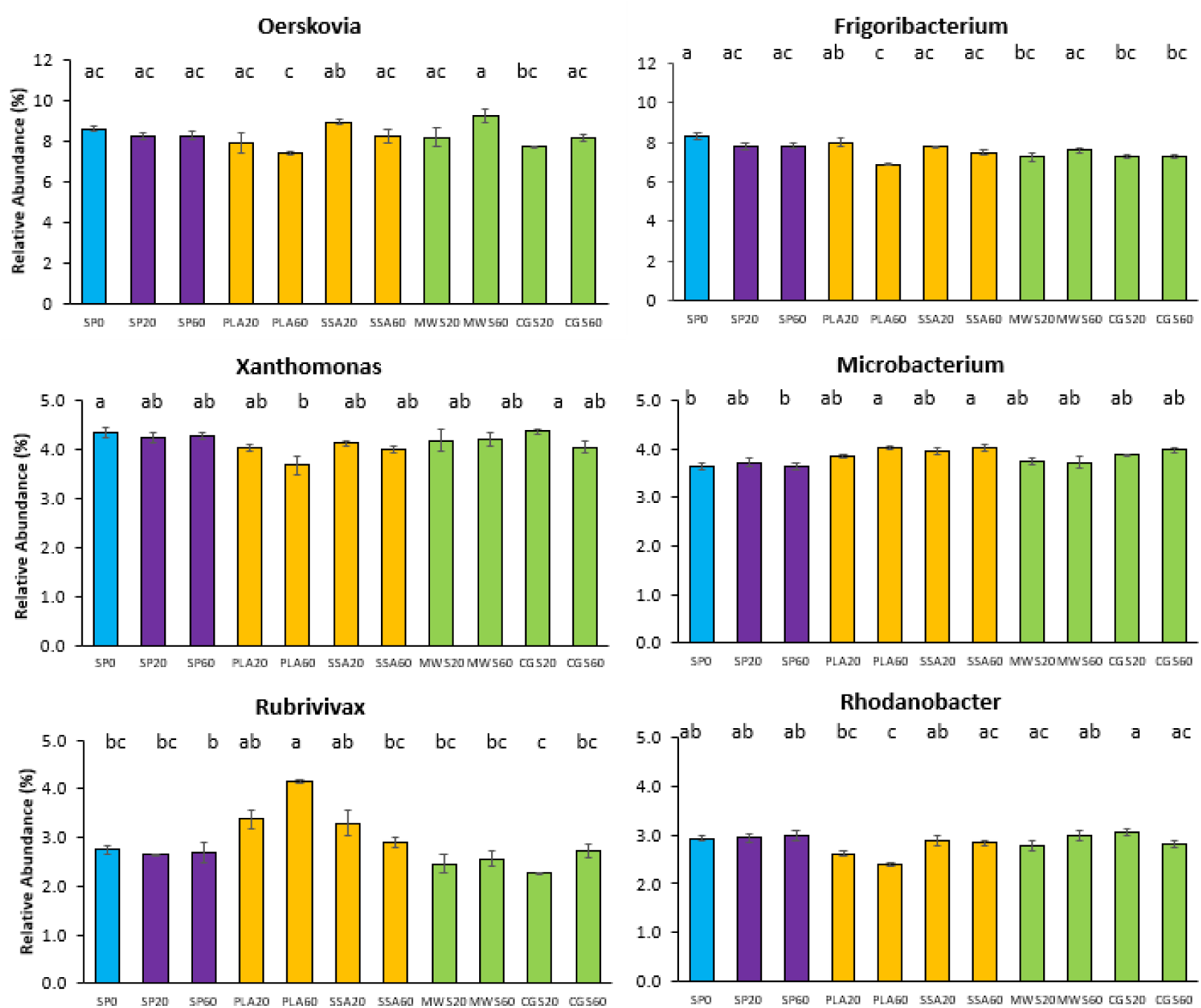
Relative abundance (%) bar plots of top 10 genera with significantly different abundance of the phoD sequencing data, SP0 in blue, SP20 and SP60 in purple, significance determined via Kruskal-Wallis test and Wilcoxon post-hoc analysis with Benjamini-Hochberg correction, different letters state significant difference, n=4.

## Discussion

The influence of four different RDFs at two different P concentrations on the perennial ryegrass dry matter yield and the P cycling soil microbiota was investigated in comparison to conventional mineral superphosphate fertilizer. At the beginning, it was hypothesised, that the RDFs provide a similar P fertilization effect as conventional mineral P fertilizer, while demonstrating less of an impact on the soil microbes involved in P mobilization activities, i.e. share similarities with he no P control.

The soil microbiota plays an important role in the nutrient cycling, especially the transformation of inaccessible P compounds into plant available orthophosphate is of interest. Their role in the nutrient cycle has received more attention in the past (Richardson & Simpson, 2011). The use of biofertilizers, microbial inoculants with known N and P mobilization traits, has been under investigation for some time, and a meta-analysis conducted by Schütz and colleagues (Schütz *et al*., 2018) found the greatest effect size for P solubilizing microorganisms at a medium plant available P concentration of 25 - 35 kg ha^-1^ (Olsen-P), however AMF performed better at lower plant available P levels between 15 - 25 kg ha^-1^ and their inoculation increased plant yield by over 40%. The detrimental effects of mineral fertilizer application on the functioning of the nutrient cycling microbiota have been reported in several studies. Mozumder and Berrens (Mozumder & Berrens, 2007) conducted a meta-analysis and described a significant correlation between amount of inorganic fertilizer applied and biodiversity loss investigating data collected across several countries. They however noted that information regarding specific fertilizer effects on the soil biodiversity were lacking in the literature.

In this study, all treatments except for PLA20 showed improved *L. perenne* dry weight yields upon P fertilizer application, with one struvite (MWS60) producing significantly higher yields than all other treatments. This indicates that all RDF treatments contain plant available P or provided suitable conditions for P to become plant available. The agronomic efficiency was variable due to low overall dry matter yields but presented its highest value for the MWS20 struvite. This is in contrast to findings in a study performed by González and colleagues (González *et al*., 2021), comparing the AE of several struvites produced from municipal wastewater to mineral triple15 fertilizer (N-P-K ratio 15:15:15). They found an increased AE for the triple15 treatment after 90 days of grass cultivation compared to the struvite treatments. However, the experiment was carried out in river sand and not in soil, limiting biological activity and therefore microbial influence on P availability was lower compared to this study. Degryse and colleagues (Degryse *et al*., 2017) argued that many studies use ground struvite mixed through soil to assess its fertilizing properties instead of granules, the latter of these likely having a slower dissolution rate. They also found that struvites dissolve much faster in acidic soils, with the dissolution rate dropping sharply from 0.43 mg per day to <0.05 mg per day for alkaline soils. In this study, the large CGS struvite granules were partially broken down, while the MWS struvite had a small particle diameter (not defined). This characteristic and the use of a rather acidic soil for this study, could have caused an improved P release from struvite RDFs, as also reflected in higher dry matter yields.

P mobilization from phosphonate and phytate showed no significant fertilizer response, although some RDFs (PLA60, SSA60 and CGS60) displayed higher average MPNs for PAA utilization than the mineral fertilizer SP60. Furthermore, P solubilization was significantly affected in the SP60 treatment. This indicates a potential negative impact of high P mineral fertilizer on the P cycling microbial community. Similarly, Fox and colleagues (2016) have reported significant higher CFU for TCP solubilizing bacteria, as well as phosphonate and phytate utilizing MPN in *Miscanthus giganteus* biochar amended soils, although only compared to a P-free control treatment.

While potential acid phosphomonoesterase activity was increased for the treatments SP60, MWS60 and CGS60 in this study, it was lower in ash RDF treatments, likely due to the liming effect of the ashes, which was detected especially at higher P ash application rates. Though there are contradicting statements about the correlation between ACP and ALP activity and soil pH (Pang & Kolenko, 1986), it is generally assumed that ACP activity is higher at low soil pH and ALP activity at high soil pH levels (Herbien & Neal, 1990, Nannipieri *et al*., 2011). However, next to soil pH, other soil properties such as soil organic matter, moisture, clay and silt, total N, isotopically-exchangeable P and extractable Mg content were significantly correlated with the intensity of phosphatase activity, according to a study conducted by Harrison (Harrison, 1983). Contrasting to findings in this study, Saha and colleagues (Saha *et al*., 2008) found an increased ACP activity in soil fertilized with compost after three years of wheat/kidney bean/corn and okra crop rotation, while inorganic NPK fertilizer treated soil demonstrated low activity. Interestingly, the ACP results in that study, investigating the impact of organic compost application on phosphatase activity, ranged between around 700 - 1500 µg p-NP g^-1^ soil, while ALP activities were roughly ten-fold lower, similar to what had been recorded in this current study under RDF fertilization. This is likely due to the acidity of the soil, where ACP activity prevails. On a molecular level however, *phoD* copy numbers were approximately thousand-fold higher abundant than *phoC*. This indicates that the high ACP activity appears to be rather linked to plant root exudation activity than direct production of the enzyme by soil microbes (Robinson *et al*., 2012).

Bacterial community structure sequencing analysis revealed differences in abundance upon different P fertilizer application in this study. *Acidobacteria* is a broadly abundant and highly phylogenetically diverse phylum that has been mostly investigated via cultivation independent methods (Kielak *et al*., 2016). In the current study, a moderate trend of higher *Acidobacteria* relative abundance in the RDF ash treatments was noted, which is in contrast to findings by Jones and colleagues (Jones *et al*., 2009), who reported that the abundance of *Acidobacteria* was strongly regulated by soil pH and its abundance is commonly higher at low soil pH levels. Alternatively, significant positive correlation between *Acidobacteria* abundance and soil organic matter and Al content was found (Navarrete *et al*., 2013). Both ash RDFs contained high amounts of Al and PLA also contained mentionable amounts of total carbon, both of which could affect *Acidobacteria* abundance positively. Significantly positive correlations have been reported for relative abundance of *Patescibacteria* with plant available P (Li *et al*., 2021, Song *et al*., 2021). This phylum was significantly higher for two struvite as well as one ash RDF in the current study, and these treatments also exhibited high biomass yield and in terms of the CGS60 treatment, a high Pav was also confirmed. Archaeal *Euryarchaeota* seemed to thrive on MWS20 struvite, while its abundance was lower for all other RDFs in the current study. Several genera of this phylum have been mentioned in the literature regarding their plant-growth promoting characteristics (Verma *et al*., 2017, Shrivastava *et al*., 2021), but detailed research remains elusive (Yadav *et al*., 2017). The phylum *Halanaerobiaeota* was predominantly found in studies assessing compost, biochar and manure. Li and colleagues (Li *et al*., 2022) reported *Halanaerobiaeota* as one of the dominant phyla present in chicken manure composting in combination with wood-derived biochar, Xie and colleagues (Xie *et al*., 2021) noted an increase in relative abundance of *Halanaerobiaeota* in the thermophilic phase of composting, and another study detected higher relative abundances of *Halanaerobiaeota* in biochar supplemented anaerobic digestates (Heitkamp *et al*., 2021). These findings lead to the assumption that *Halanaerobiaeota.* might be sensitive towards mineral P inputs, as the RDFs as well as the SP treatment contain forms of mineral P. On genus level then, *Nocardioides*, *Conexibacter* and *Clostridium sensu stricto 13* were negatively affected by the PLA60 treatment compared to all other high P fertilization treatments in this study. All three genera have been reported to participate in P cycling activities (Deng *et al*., 2021, Ma *et al*., 2021, Wang *et al*., 2021).

Similar to the bacterial 16S rRNA phylogenetic analysis, the evaluation of the *phoD* harbouring community did not reveal significant shifts in a CCA, although some trends were observed in the form of a separation of MWS60 on the first, and SSA60 on the second axis.

Furthermore, alpha diversity of the *phoD* harbouring community was not significantly different. Differential abundant genera *Oerskovia*, *Frigoribacterium*, *Xanthomonas* and *Rhodanobacter* were significantly lower abundant in PLA60, potentially impacting bacterial P mobilization originating from alkaline phosphomonoesterase. Of the most abundant genera identified in this current study, *Rhodanobacter*, *Streptomyces* and *Microbacterium* were also detected in the rhizosheath (sampling occurred <1 mm from the root) in a study assessing struvite fertilizer application on rhizosphere dynamics via 16S sequencing (Robles-Aguilar *et al*., 2020).

In conclusion, struvite RDFs demonstrated improved ryegrass yields via higher P availability while simultaneously maintaining P mobilization activities in soil. Mineral P application via superphosphate had detrimental effects on TCP solubilizing bacteria. These findings partially support the hypotheses of improved P availability und an RDF regime, while having less impact on the P cycling microbiota. Ash RDFs however showed negative impacts on abundance of P mobilizing genera, a more variable performance, likely due to their different feedstocks, containing substances that could constrain soil microbes. Further research has to be conducted on RDF use in grasslands, especially to determine long-term effects of the rather slow-release dynamics of these novel fertilizers on the soil P mobilizing community and long-term P plant availability.

## Acknowledgements

We would like to acknowledge Interreg North West Europe ReNu2Farm (NWE601) for financing this study.

